# The actin networks of chytrid fungi reveal evolutionary loss of cytoskeletal complexity in the fungal kingdom

**DOI:** 10.1101/2020.06.12.142943

**Authors:** Sarah M. Prostak, Kristyn A. Robinson, Margaret A. Titus, Lillian K. Fritz-Laylin

## Abstract

Cells from across the eukaryotic tree use actin polymers and a number of conserved regulators for a wide variety of functions including endocytosis, cytokinesis, and cell migration. Despite this conservation, the actin cytoskeleton has undergone significant evolution and diversification, highlighted by the differences in the actin cytoskeletal networks of mammalian cells and yeast. Chytrid fungi diverged before the emergence of the Dikarya (multicellular fungi and yeast), and therefore provide a unique opportunity to study the evolution of the actin cytoskeleton. Chytrids have two life stages: zoospore cells that can swim with a flagellum, and sessile sporangial cells that, like multicellular fungi, are encased in a chitinous cell wall. Here we show that zoospores of the amphibian-killing chytrid *Batrachochytrium dendrobatidis (Bd)* build dynamic actin structures that resemble those of animal cells, including pseudopods, an actin cortex, and filopodia-like actin spikes. In contrast, *Bd* sporangia assemble actin patches similar to those of yeast, as well as perinuclear actin shells. Our identification of actin cytoskeletal elements in the genomes of five species of chytrid fungi indicate that these actin structures are controlled by both fungal-specific components as well as actin regulators and myosin motors found in animals but not other fungal lineages. The use of specific small molecule inhibitors indicate that nearly all of *Bd*’s actin structures are dynamic and use distinct nucleators: while pseudopods and actin patches are Arp2/3-dependent, the actin cortex appears formin-dependent, and actin spikes require both nucleators. The presence of animal- and yeast-like actin cytoskeletal components in the genome combined with the intermediate actin phenotypes in *Bd* suggests that the simplicity of the yeast cytoskeleton may be due to evolutionary loss.

## INTRODUCTION

Actin participates in nearly every essential eukaryotic cell function, including endocytosis, intracellular trafficking, and cell migration, as well as in cytokinesis in many species (reviewed in (Velle and Fritz-Laylin 2019). Eukaryotic cells employ a sophisticated network of actin regulatory proteins to spatially and temporally control these diverse functions. How these complex actin regulatory networks evolved and diversified remain key questions in both evolutionary and cell biology. Here we use chytrids—early-diverging fungi that still share important features of animal cells lost in yeast and other fungi (Hodges et al. 2010; Medina et al. 2016)—as a system to explore the evolution of the actin cytoskeleton. Using a combination of genomics and fluorescence microscopy, we show that chytrid fungi have an actin cytoskeleton that combines features of animal cells as well as yeast.

Actin polymerization is largely regulated by controlling the initiation of new actin polymers—a process called “actin nucleation.” Actin is nucleated by two main systems: the Arp2/3 complex or formin family proteins, both of which were likely present in the last common eukaryotic ancestor (Beltzner and Pollard 2004; Chalkia et al. 2008; Veltman and Insall 2010; Kollmar et al. 2012; Pruyne 2017; Velle and Fritz-Laylin 2019). While the Arp2/3 complex primarily builds actin branches along the side of existing actin filaments (Mullins et al. 1998; Svitkina and Borisy 1999), formins assemble unbranched filament networks through processive addition of actin monomers by their Formin Homology 2 (FH2) domains (Pruyne et al. 2002; Sagot et al. 2002; Kovar et al. 2003; Zigmond et al. 2003; Goode and Eck 2007; Breitsprecher and Goode 2013).

Animal cells use the Arp2/3 complex and formins to build a wide variety of dynamic actin structures. These structures include diverse membrane protrusions used for movement (Fritz-Laylin et al. 2018), from broad pseudopods and lamellipodia that are filled with branched-actin networks (Svitkina and Borisy 1999; Fritz-Laylin, Riel-Mehan, et al. 2017; Fritz-Laylin, Lord, et al. 2017; Blanchoin et al. 2014), to finger-like filopodia that are packed with linear actin bundles (Gupton and Gertler 2007; Mellor 2010; Gallop 2020). Dynamic actin networks and their associated motors also mediate membrane invagination during endocytosis (Mooren et al. 2012; Lu et al. 2016; Carlsson 2018) as well as the construction of the cytokinetic ring (Cheffings et al. 2016; Mangione and Gould 2019). Many of these actin structures often assemble in proximity to the “actin cortex,” a 200 nm thick actin shell that lies just below and supports the plasma membrane (Clark et al. 2013; Haase and Pelling 2013).

Budding and fission yeast, in contrast to animal cells, each have simplified actin networks that consist of three main structures: actin patches that are sites of endocytosis and deposition of cell wall material (Kaksonen et al. 2003; Young et al. 2004; Kaksonen et al. 2005; Sirotkin et al. 2010); actin cables for vesicle trafficking and establishing cell polarity (Moseley and Goode 2006; Drake and Vavylonis 2010; Bergs et al. 2016); and the cytokinetic actin ring (Cheffings et al. 2016; Mangione and Gould 2019). This simplification is echoed by a streamlined actin regulatory system that is missing key actin regulators important for human health (Fritz-Laylin, Lord, et al. 2017), particularly the SCAR/WAVE complex which helps drive cell migration involved in normal development of mouse embryos, and in metastasis and tissue invasion in tumor models (Rakeman and Anderson 2006; Kurisu and Takenawa 2009; Burianek and Soderling 2013; Moazzam et al. 2015). The wide gap between mammalian and yeast actin biology makes it difficult to know which rules of yeast actin regulation apply to human cell biology. Bridging this gap would allow us to apply our deep understanding of simplified yeast actin networks to human cells.

To help us bridge the gap between the simplified actin networks of yeast and the dizzyingly complex actin networks of human cells, we turned to chytrid fungi. Chytrids play key roles in aquatic and terrestrial habitats (Richards et al. 2012; Sime-Ngando 2012; Kagami et al. 2014; Grossart et al. 2016), including *Batrachochytrium dendrobatidis* (*Bd*), an amphibian pathogen that is decimating frog populations across the globe (Berger et al. 1998; Longcore et al. 1999; Lips 2016; O’Hanlon et al. 2018). Also known as zoosporic fungi, the >1000 known species of chytrids comprise at least three fungal phyla: Cryptomycota, Chytridiomycota, and Blastocladiomycota (James et al. 2006). Chytrids diverged from a common fungal ancestor before the diversification of the Dikarya (James et al. 2006), the group of fungi that includes multicellular mushrooms and sac fungi, as well as unicellular yeasts that descended from multicellular fungi (**Fig. 1a**). These phylogenetic relationships position chytrids as an evolutionary Rosetta Stone with which we can map the simplified actin features of yeast to those of animals.

**Figure 1.**
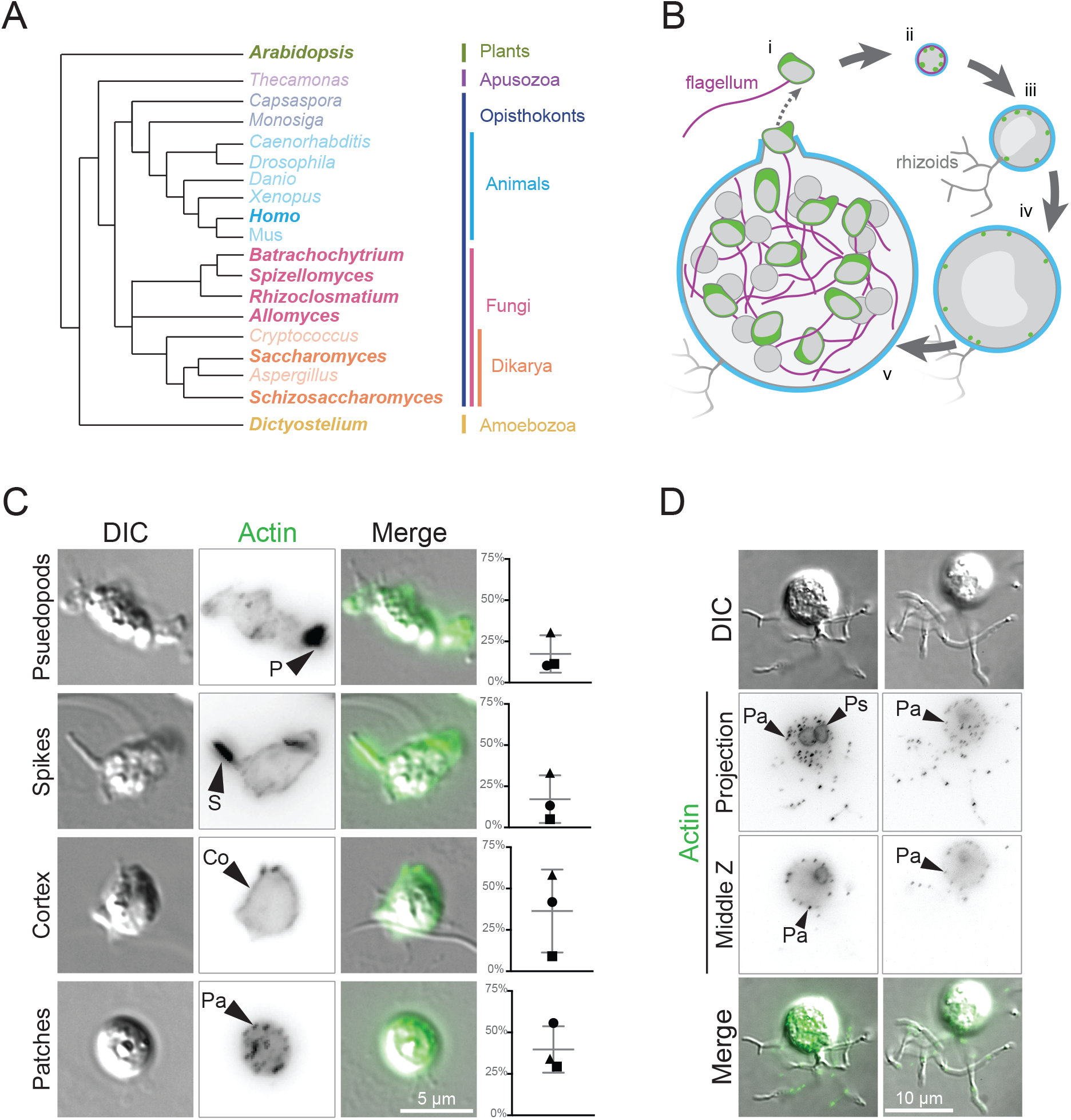
The chytrid fungus Batrachochytrium dendrobatidis is an early branching fungus with an archetypal chytrid life cycle and animal-like and fungal-like actin structures. (A) This cladogram shows the relationships between representative genera of major eukaryotic groups. Chytrids are represented by *Batrachochytrium*, *Spizellomyces*, *Rhizoclosmatium*, and *Allomyces* (magenta), diverging before the diversification of Dikarya (orange), and are in a sister clade to animals (cyan). Genera that are bolded were used for the majority of the homologous sequence analyses in this paper. (B) *In vitro* life cycle of *Batrachochytrium dendrobatidis (Bd)*. Bd has a motile stage known as a zoospore (i) with a flagellum made of microtubules (magenta), no cell wall, and can crawl using actin (green) based protrusions. Zoospores encyst and build a cell wall (cyan), this stage is referred to as a germling (ii). The germling grows in size, becoming a sporangium. Sporangia develop hyphal-like structures called rhizoids used for nutrient uptake and undergo synchronous rounds of mitosis (iii-iv) before cellularization and release of the next generation of zoospores (v). This life cycle takes approximately three days in laboratory culture conditions. (C) Representative examples of zoospores (DIC: grey) and the phalloidin stained actin structures at this stage (inverted, black), with an overlay of the two (actin, green). Actin structures in *Bd* zoospores are: actin-filled pseudopods (P), actin-filled spikes (S), cortical actin (Co), and actin patches (Pa). Graphs on the right indicate the raw percent of cells with the phenotype in 3 independent experiments, shapes here match the shapes for the replicates in Fig. 6. (D) Representative examples of *Bd* sporangia (DIC: grey) and the phalloidin stained actin structures at this stage (inverted, black; both a max intensity z projection and a single slice, with an overlay of the DIC and fluorescence (actin, green). Actin structures in sporangia are: actin patches (Pa), and perinuclear actin shells (Ps).

*Bd*, like other chytrids, has two developmental stages: motile “zoospore” cells that lack a cell-wall and swim with a flagellum, and non-motile sporangia that grow and produce new zoospores (**Fig. 1b**; (Berger et al. 2005; Longcore et al. 1999). We recently showed that zoospores can also crawl across surfaces using actin-filled, Arp2/3-dependent pseudopods (Fritz-Laylin, Lord, et al. 2017). Other than these pseudopods, actin’s role in any developmental stage of *Bd* has not been studied. To fill this knowledge gap, we identified homologs of key actin regulatory proteins and myosin motors in multiple species of chytrids and related fungi. We also identified new actin structures in zoospores and sporangia, and tested the requirements of Arp2/3 and formin family proteins for their assembly. We find that both the regulatory networks and actin structures of *Bd* are intermediate in complexity between animals and Dikarya, suggesting that the streamlined actin networks of common model fungi are a result of secondary evolutionary loss of actin network components.

## RESULTS

### Bd has developmentally distinct actin phenotypes that resemble animal and fungal cells

To determine how *Bd* may use actin during its life cycle, we stained *Bd* cells with fluorescent phalloidin that labels polymerized actin (Estes et al. 1981; Fritz-Laylin, Lord, et al. 2017). Staining of *Bd* zoospores revealed four easily distinguishable actin structures found in various combinations (**Fig. 1c**, **Fig. S1**, **Fig. S2**): (**1**) pseudopods that are roughly 1-2 μm across; (**2**) filopodia-like actin “spikes” that have not been previously described in *Bd*, which we define as thin, actin-filled protrusions <1 μm across and ≥1μm long; (**3**) cortical actin localized to more than half of the cell edge; and (**4**) <1 μm diameter actin puncta. We call these last structures “actin patches” due to their visual similarity to yeast actin patches (Kaksonen et al. 2003; Young et al. 2004; Kaksonen et al. 2005; Sirotkin et al. 2010) as well as their localization at the cell periphery (**Fig. 1d**). The percent of zoospores with each actin structure is variable from experiment-to-experiment (**Fig. 1c**), likely due to slight differences in developmental staging between biological replicates.

We also stained *Bd* sporangia—large multinucleated cells that undergo mitosis and expansive cell growth while encased in a cell wall (**Fig. 1biii-v**). *Bd* sporangia stained for polymerized actin show two types of actin structures: perinuclear shells, defined by intense F-actin staining around each nucleus, similar to those observed in *Spizellomyces punctatus* (Medina et al. 2020); and actin patches (**Fig. 1d**). Both structures were present in 100% of observed sporangia.

### The actin regulatory network of chytrids resembles that of both animals and Dikarya

To explore how *Bd*’s animal- and yeast-like actin structures may be built and regulated, we performed BLAST searches for actin and across five chytrid species: *Bd*, *Batrachochytrium salamandrivorans* (*Bs*), *Spizellomyces punctatus* (*Sp*), *Rhizoclosmatium globosum* (*Rg*), and *Allomyces macrogynus* (*Am*) (**Fig. 2**, **Fig. S3 Fig. S4**, **Table S1**), and compared these to homologs from humans, *Arabidopsis thaliana*, *Dictyostelium discoideum*, *Schizosaccharomyces pombe*, and *Saccharomyces cerevisiae*. These analyses, summarized here, revealed that chytrids share a variety of actin regulators with animals that have been lost in the Dikarya. For the full description of these results, see **Supplemental Text 1**.

**Figure 2.**
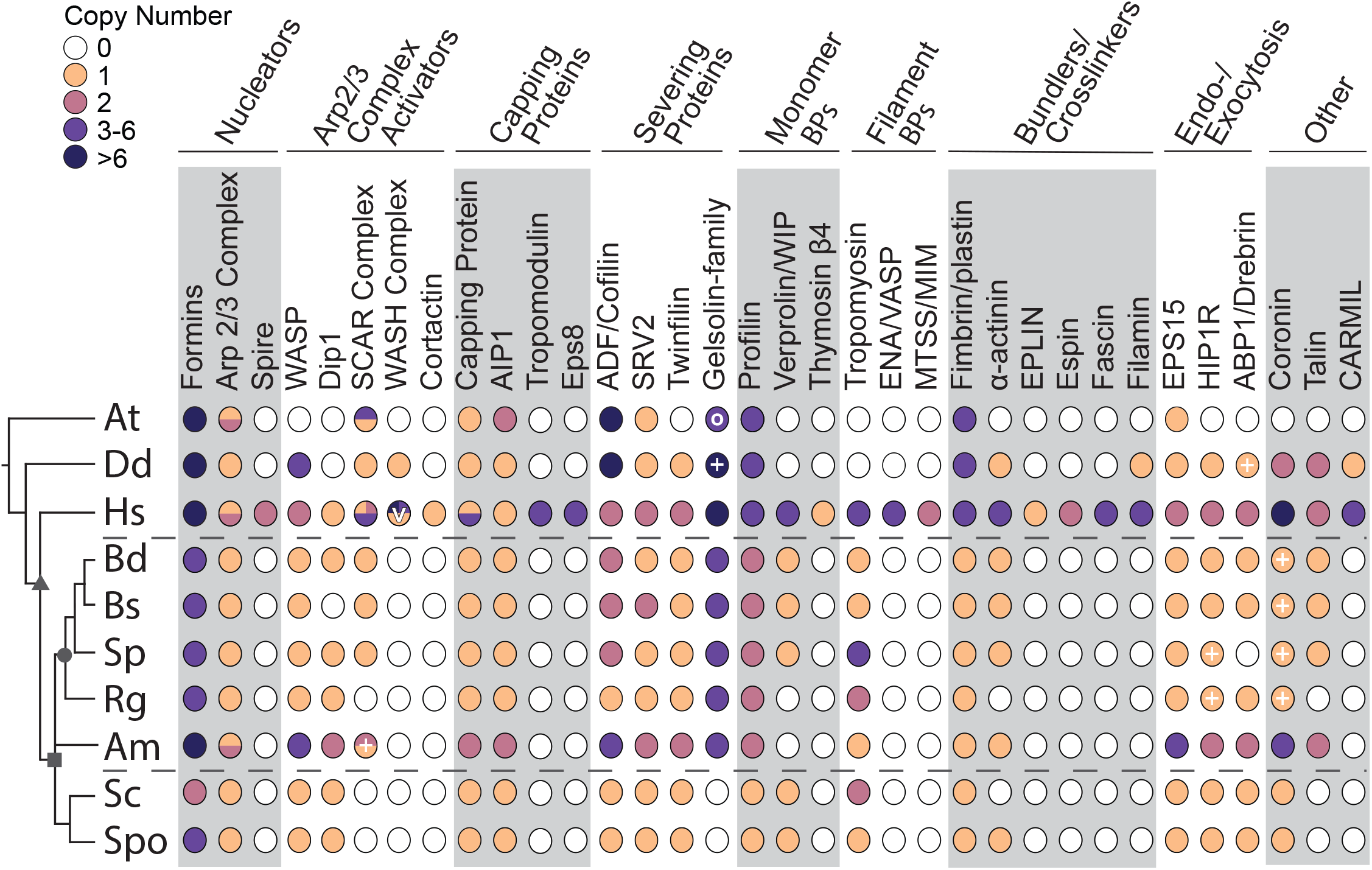
Chytrid actin regulatory protein networks are intermediate to those of animals and Dikarya. The distribution of actin regulatory proteins across taxa. Color-filled circles indicate the presence of clear homologs found, with the number of homologs for each protein in each species shown in the colors specified in the key. Unfilled circles indicate that no homolog was detected in that species. Circles with multiple colors indicate complexes with different copy numbers for multiple complex members. Circles for Capping Protein represent the copy number for both α and β subunits. Dashed lines mark the chytrids. Symbols on the tree represent: opisthokonts (triangle); fungi (square); chytridiomycota (circle). Symbols in the circles represent: V, copy number of WASH varies individually, as many WASH genes are subtelomeric (Linardopoulou et al. 2007). O, Arabidopsis has 5 villin-like genes and an additional gelsolin-domain containing protein, none of which are phylogenetically related to metazoan gelsolin/villin family members (Ghoshdastider et al. 2013). +, See **Supplemental Table 1** for details and additional potential homologs with caveates. *At, Arabidopsis thaliana; Dd, Dictyostelium discoideum; Hs, Homo sapiens; Bd, Batrachochytrium dendrobatidis; Bs, Batrachochytrium salamandrivorans; Sp, Spizellomyces punctatus; Rg, Rhizoclosmatium globosum; Am, Allomyces macrogynus; Sc, Saccharomyces cerevisiae; Spo, Schizosaccharomyces pombe*.

Cells use two major classes of actin nucleators to stimulate actin nucleation: formin family proteins that nucleate unbranched actin filaments *de novo (Pruyne et al. 2002; Sagot et al. 2002; Kovar et al. 2003)* and the Arp2/3 complex, which typically nucleates a new actin filament on the side of existing filaments to form branched actin networks (Mullins et al. 1998; Svitkina and Borisy 1999). We find that each chytrid species has at least 4 formins with a diversity of domain organizations that we describe in detail below. All chytrid species also have at least one copy of each of the 7 members of the Arp2/3 complex (**Fig. S4**), as well as a variety of Arp2/3 activators. Some Arp2/3 activators are conserved in humans, chytrids, and Dikarya, such as WASP, and WISH/Dip1/SPIN90 (with the exception of *Bs*) (**Fig. S4**). The SCAR/WAVE complex, in contrast, is present in humans and most chytrids (SCAR/WAVE complex is missing from *Rg*, **Fig. S4**), but has been lost from the Dikarya (Veltman and Insall 2010; Kollmar et al.2012; Fritz-Laylin, Lord, et al. 2017). The tight association between the SCAR/WAVE complex and cell migration suggests its retention correlates with cell migration (Fritz-Laylin, Lord, et al. 2017), suggesting *Rg* may not crawl.

Actin is also subject to negative regulation, particularly by capping proteins that prevent further filament elongation and severing proteins that cut existing filaments. Many of these proteins are conserved in humans, chytrids, and Dikarya, such as primary capping proteins CapZ and AIP1, primary severing proteins Colfilin and Twinfilin, as well as the severing catalyst SRV2 (**Fig. 2**). The only protein/protein family to be conserved in animals and chytrids, but not in other fungi, is the gelsolin/villin family of proteins (**Fig. 2**). The conservation of these negative regulators indicates that *Bd*’s actin structures are likely highly dynamic.

We also identified a number of actin binding proteins conserved in humans, most chytrids, as well as Dikarya, including: Verprolin/WIP; Tropomyosin; Fimbrin/plastin; α-actinin; endo-/exocytosis proteins EPS15/Ede1, HIP1R/Sla2, and drebrin-like/ABP1 (**Fig. 2**). Interestingly, we find that talin is not found in Dikarya, but is conserved in most chytrids (with the exception of *Rg*, **Fig. 2**). Talin links adhesion receptors to the actin cytoskeleton in crawling cells (Niewöhner et al. 1997; Priddle et al. 1998; Brown et al. 2002; Cram et al. 2003) and may be used during zoospore crawling, a hypothesis consistent with *Rg*’s lack of both talin and the SCAR/WAVE complex.

Taken together, we find chytrid genomes encode a network of actin cytoskeletal regulators that is intermediate to the networks of animals and fungi, including SCAR/WAVE complex, gelsolin/villin family proteins, and talin, which are not found in Dikarya. For an in depth overview of all of the actin cytoskeletal components we identified, see **Supplementary Text 1**.

### Chytrids have typical fungal myosins as well as MyTH-FERM myosins

The myosin superfamily of actin-based motors plays diverse roles in cells, including providing contractile forces during cell migration and cytokinesis, powering organelle transport, driving endocytosis and building or maintaining actin-based structures such as filopodia. We searched for myosins in the same five species of chytrids and found the same core group of myosins seen in the majority of fungi - Myo1, Myo2, Myo5 and Myo17 (**Fig. S5**) as well as Myo22 (**Fig. S5**, **Supplementary Text 1**) (Odronitz and Kollmar 2007; Kollmar and Mühlhausen 2017):

Myo1s are ancient, widely expressed myosins that link membranes to the actin cytoskeleton (McIntosh and Ostap 2016; Kollmar and Mühlhausen 2017). The budding and fission yeast Myo1s recruit activators of the Arp2/3 complex to actin patches to drive internalization of endocytic vesicles (Giblin et al. 2011). Each chytrid species has one or two Myo1s, which we predict serve the same function and localize to the cortical actin patches observed in *Bd* and *Sp* zoospores and sporangia (**Fig 1**; (Medina et al. 2020).

In contrast to the ubiquity of Myo1s, Myo2s are found mainly in Amoebozoans and Opisthokonts where they are an essential component of the cytokinetic contractile ring, generating forces necessary for the scission of two daughter cells during the final steps of mitosis (West-Foyle and Robinson 2012). Myo2s also play key roles in cell migration of animal cells and Amoebozoa, where they drive the retrograde flow of the actin network and generate cell polarity in migrating cells by contracting the actin network at the cell rear (Aguilar-Cuenca et al. 2014). Chytrid fungi have a single Myo2 (**Fig. S5**) that may play roles in zoospore crawling as well as cellularization (**Fig. 1**).

Myosins also play key roles in intracellular transport, particularly Myo5s, which are present in Amoebozoa, Apusozoa, and Opisthokonts where they serve as actin-based transporters and localize cargo to the actin cortex (Titus 2018). Hyphal fungi use microtubules for long-distance transport and in these species Myo5s collaborate with kinesins. Many fungi have two Myo5s with distinct cellular functions. For example, one Myo5 of *S. cerevisiae* is required for organelle inheritance, polarized budding and mitotic spindle orientation while the second one is critical for polarized localization of cell fate determinants (Matsui 2003). All five chytrid species contain at least one Myo5 that likely plays critical roles in intracellular transport, aiding in organelle segregation during division or targeting vesicles to sites of polarized growth.

Chytrids also have Myo17s. These unusual chimeric fungal myosins have a core motor domain arm fused to a chitin synthase 2 (Ch2) domain (Kollmar and Mühlhausen 2017). Transmembrane domains orient the protein so that upon exocytosis the chitin synthase enzyme is positioned toward the outer cell wall. Myo17s are essential for cell wall integrity and virulence in several pathogenic fungi (Takeshita et al. 2006; Gandía et al. 2014), and interestingly, a highly similar myosin is also found in molluscs (Weiss et al. 2006). Like other fungi, chytrids have multiple Myo17 family members that are likely used for targeted cell wall synthesis during different growth stages.

Although all chytrid species analyzed have at least one copy of Myo1, 2, 5, and 17, only some chytrid species appear to have retained Myo22. These myosins have one or two MyTH-FERM domains in the C-terminal tail region (Kollmar and Mühlhausen 2017) and are largely associated with the assembly and function of cellular protrusions composed of parallel bundles of actin, such as the filopodia of *Dictyostelium* and animal cells (Tuxworth et al. 2001; Petersen et al. 2016; Weck et al. 2017). Although a subset of chytrid fungi including *Rg* and *Sp* have a single Myo22, neither *Am*, *Bd* nor *Bs* have a Myo22. We have not been able to detect these myosins in any other fungal species outside of the chytrids, suggesting that this myosin was lost early in fungal evolution. The potential function of Myo22 in *Rg* and *Sp* is unclear at present but it may contribute to the formation of a distinctive actin-based structure specific to these species.

### Chytrid formins resemble those of fungi, animals, and plants, including DAAM and other diaphanous-related formins lost among the Dikarya

One of the largest families of actin regulators in chytrids is the formin family proteins (**Fig. 2**). Given the importance of these actin nucleators to animal and fungal cell biology—formins nucleate the actin networks used for cytokinesis, cell movement, filopodia, and vesicle trafficking (Breitsprecher and Goode 2013)—we next sought to explore the diversity and evolution of formin protein sequences in chytrids. Although formins are defined by an FH2 domain that nucleates actin polymerization (Pruyne et al. 2002; Sagot et al. 2002; Kovar et al. 2003), the biological function of each formin is heavily influenced by additional and highly variable protein domains that regulate its function and localization. Humans, for example, have 15 formins that are divided into seven main groups, four of which have a similar domain organization, known as the diaphanous-related formins, while the other remaining three have their own unique domain organizations (Breitsprecher and Goode 2013). Although yeast formins share some similarity in domain organization to metazoan formins, yeast have far fewer formin genes—2 in *S. cerevisiae* and 3 in *S. pombe (Chalkia et al. 2008; Breitsprecher and Goode 2013)*.

The complexity of the formin protein family (**Fig. 3**, **Fig. 4**) prompted us to further investigate these proteins in chytrids by first determining their domain organizations. We identified the conserved domains of each chytrid formin using the Pfam database, and manually inspected protein sequences for the presence of FH1 domains and diaphanous autoregulatory domains. Here we describe the two most common domain organizations: Diaphanous-like formins and PTEN-like formins. For additional domain information see **Supplementary Text 1**.

**Figure 3.**
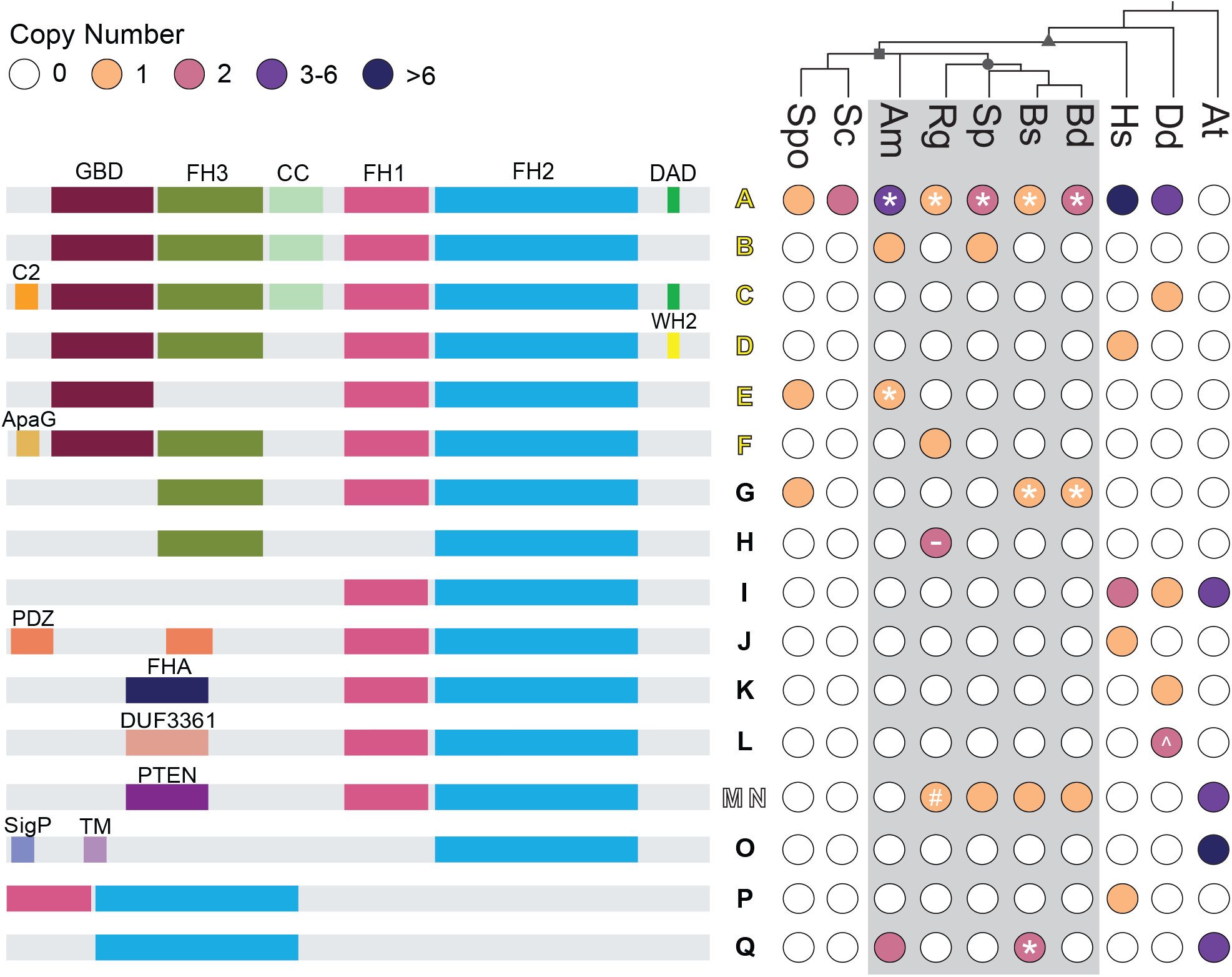
Chytrid formins share similar domain architectures to those of animals, Dikarya, and plants. The distribution of given formin domain architectures (left, not to scale), across taxa (right). Each domain architecture is assigned a letter (middle) which is mapped onto Figure 5; yellow letters (A-F) indicate diaphanous-like architectures, white letters (M,N) indicate PTEN-domain-containing architectures (both plant and non-plant formins), black letters indicate architectures which did not fall into either of these classes. Color-filled circles (right) indicate the presence of at least one formin with the given domain architecture in that species, with color indicating the number of formins. Unfilled circles indicate that no formin with the indicated domain architecture was found in the given species. Symbols on the tree represent: opisthokonts (triangle); fungi (square); chytridiomycota (circle). Symbols in the circles represent: *, for at least one formin sequence in the indicated species, the DAD domain does not perfectly fit the consensus motif, but could potentially function as an autoregulatory domain. -, for at least one formin sequence, little to no sequence is present after the FH2 domain. #, although no region of this protein met the formal definition of an FH1 domain, a proline rich region (4 polyproline stretches: 8 prolines/14 amino acids; 4/5; 4/6; and 7/11; total of 22 prolines over 197 amino acids) is found N-terminal to the FH2 domain in this protein (Genbank: ORY46833.1). ^, Dictyostelium formin ForC has no polyproline stretches and therefore no FH1 domain. *At, Arabidopsis thaliana; Dd, Dictyostelium discoideum; Hs, Homo sapiens; Bd, Batrachochytrium dendrobatidis; Bs, Batrachochytrium salamandrivorans; Sp, Spizellomyces punctatus; Rg, Rhizoclosmatium globosum; Am, Allomyces macrogynus; Sc, Saccharomyces cerevisiae; Spo, Schizosaccharomyces pombe*.

**Figure 4.**
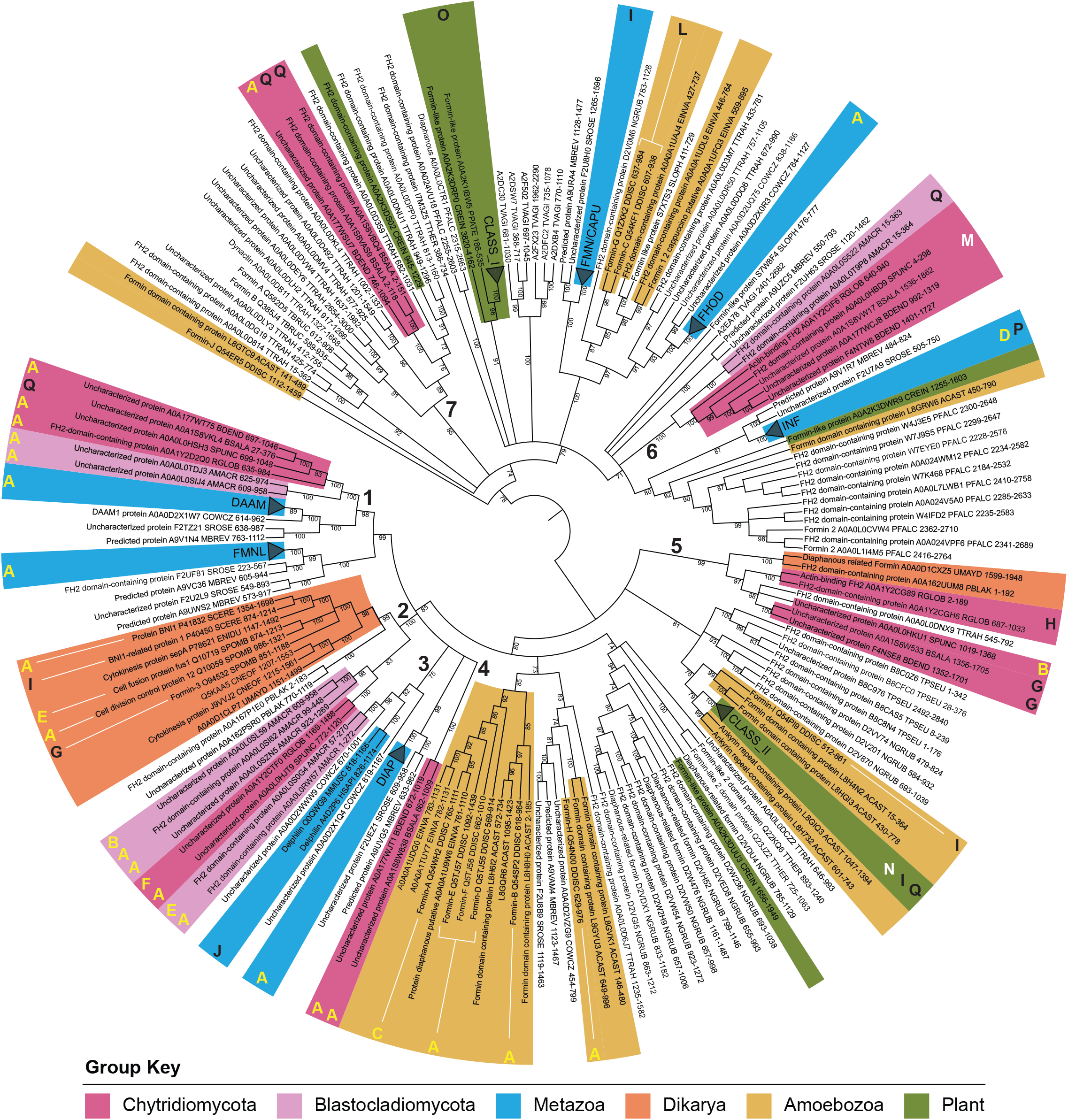
Chytrids have animal-like formins related to DAAM as well as other diaphanous-related formins. A maximum likelihood consensus tree was inferred using the FH2 domains of 259 formin proteins, rooted at the midpoint, with bootstrap values as shown, and nodes with <70% bootstrap support collapsed to polytomies. Metazoan clades and *Arabidopsis thaliana* clades were collapsed and named according to their formin group, except for Delphilin. Taxa of interest are colored according to the key, bold numbers indicate chytrid-containing clades. The bold letters around the outside of the tree correspond to the domain architectures in **Figure 3**; yellow letters indicate diaphanous-like architectures, white letters indicate PTEN-domain-containing architectures, black letters indicate architectures which did not fall into either of these classes. Protein names, Uniprot accession numbers, Full species name, Uniprot 5-letter species codes, position of the FH2 domain, and additional details can be found in **Supplemental Data File 5**.

The diaphanous-related formins have an N-terminal GTPase-binding domain (GBD) that binds to Rho GTPases to release the inhibitory interaction between the N-terminal inhibitory domain (FH3/DID) and the C-terminal diaphanous auto-regulatory domain (DAD) (Rivero et al. 2005; Breitsprecher and Goode 2013). Each chytrid species contains at least one formin with a domain architecture that resembles that of diaphanous-related formins; containing a GTPase-binding domain, an FH3/DID domain, a coiled coil region, and an FH1 domain all N-terminal to the FH2 domain (**Fig. 3**). As for the C-terminal diaphanous autoregulatory domain, only three proteins had the consensus sequence (MDXLLXXL; (Alberts 2001; Higgs 2005)), while the remaining proteins had similar sequences which may be functional if compensatory substitutions occurred in the FH3/DID (**Fig. 3**, **Data S2**). For detailed information on additional chytrid diaphanous-like formins which stray slightly from the canonical organization see **Supplementary Text 1**.

Each chytrid, except *Am*, has a formin containing a PTEN/PTEN-like domain N-terminal to the FH2 domain (**Fig. 3**). PTEN-formins are best known for their role in plants where they mediate membrane-localization by binding phospholipids (Grunt et al. 2008; Pruyne 2017). *Arabidopsis*, for example, has 10 Class II formins, 4 of which have N-terminal PTEN-like domains that localize these formins to membranes (Deeks et al. 2002; Cvrcková et al. 2004; Grunt et al. 2008; Breitsprecher and Goode 2013). The PTEN-like domain of class II formins has also been shown to mediate formin-microtubule interactions (Kollárová et al. 2020). Despite having similar domain organization, our phylogenetic analysis of FH2 domains (see below) suggests that these chytrid formins are not related to Class II plant formins and may represent a secondary gain of a PTEN/PTEN-like domain (Pruyne 2017).

We next sought to determine whether the Diaphanous-like and PTEN-like formins were homologous to formins with similar domain structures from animals and plants. Formins are a diverse family in terms of domain organization, and it would be tempting to assume that structurally similar formins are evolutionarily related. However, structural similarity does not necessarily indicate relatedness, as the formin family has likely undergone gene duplication, divergence, and domain shuffling throughout its history (Chalkia et al. 2008; Pruyne 2017). A great example of this are the fungal formins, as most fungal formins are organized similarly to several human formins, but are phylogenetically distinct (Higgs and Peterson 2005; Chalkia et al. 2008; Pruyne 2017). We therefore wanted to determine how chytrid formins fit into this complex family history. We aligned 259 FH2 domain-containing sequences from 28 species across eukaryotic phyla to the Pfam FH2 domain (PF02181) full hidden Markov model, isolated the FH2-domains based on the predicted positions of the FH2 domain for *S. cerevisiae* Bni1p, and created a Maximum Likelihood phylogeny (**Fig. 4; Data S3, Data S4, Data S5**).

Our phylogeny reveals that, surprisingly, most chytrid formins do not form a neat clade with other fungal formins, but instead are scattered throughout the tree in at least seven distinct clades (**Fig. 4**), including: (**1**) DAAM-related formins with homologs in animals and their unicellular relatives; (**2**) *bni1*-type formins with homologs in other fungi; (**3**) Delphilin-related formins that have homologs in *Allomyces* but not in Chytridiomycetes; (**4**) a clade of formins found only in the parasitic chytrids *Bd* and *Bs*; (**5**) Fungi-2, a second clade of fungal formins known to be present in some fungi (Pruyne 2017); (**6**) non-plant PTEN/PTEN-like formins; and (**7**) a second *Bd* and *Bs* clade that also includes a homolog from the flagellated green algae *Chlamydomonas*. Proteins within each clade tend to share overall domain architectures. Interestingly, the chytrid PTEN/PTEN-like formins are not related to plant Class II formins (**Fig. 4**), suggesting that their similarity in domain architecture may be due to convergent evolution (Pruyne 2017).

### Bni1-type formins are associated with actin cables

Our phylogeny supports previous findings that yeast formins form their own clade separate from animal formins (Higgs and Peterson 2005; Chalkia et al. 2008; Pruyne 2017), which we will refer to as the *bni1*-type formins (**Fig. 4,** group 2). In budding yeast, *bni1p* nucleates actin cables originating in the bud while *bnr1p* nucleates actin cables primarily at the bud neck and into the mother cell (Pruyne et al. 2004; Buttery et al. 2007). However in fission yeast, each formin has distinct non-overlapping functions, and For3 is responsible for nucleating actin cables (Feierbach and Chang 2001). We find that some chytrids have at least one *bni1*-type formin, including *Sp*, whose sporangia we recently found can build actin cables (Medina et al. 2020). Other chytrids, including *Bd* and *Bs*, are missing these formins, an interesting finding given the lack of actin cables in *Bd* zoospores and sporangia (**Fig. 1**).

We therefore hypothesized that a fungal species must have a *bni1*-type formin to build actin cables. To explore this hypothesis, we combed the literature to identify chytrid species with observed actin cables and identified three chytrid species that have been stained for actin, all of which have actin cables: *Neocallimastix patriciarum (Li and Heath 1994); Orpinomyces joyonii (Li and Heath 1994)*; and *Chytriomyces hyalinus (Dee et al. 2019)*. We added the formin FH2 domains from these species to the phylogeny to determine if they had at least one *bni1*-type formin. Due to the lack of publicly available genome sequences for *N. particiarum* and *O. joyonii*, we used the genomes of species in the same genus (*Neocallimastix californiae* G1 and *Orpinomyces sp.* strain C1A). We find that all three additional cable-containing chytrid species added to the phylogeny have a *bni1*-type formin (**Fig. 5a**). We also fixed and stained *Bs* sporangia at several time points in its lifecycle. We find that *Bs*, a chytrid with no *bni1*-type formin, has no actin cables throughout its sporangial stage, unlike *Sp* sporangia, which have pronounced cables extending into the rhizoids (**Fig. 5b**). These findings are consistent with our hypothesis that *bni1*-type formins are used to build actin cables (**Fig. 5c**).

**Figure 5.**
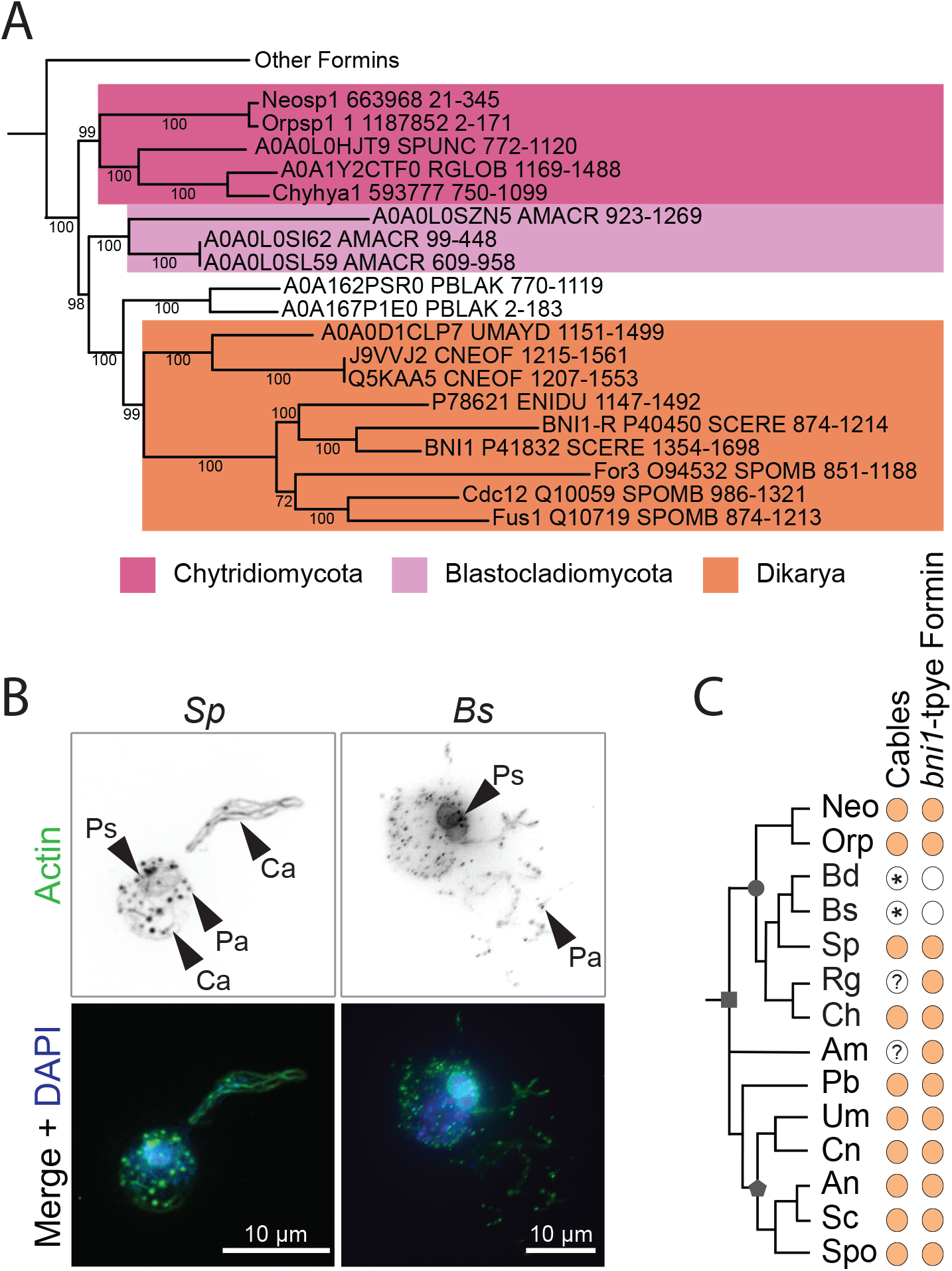
Bni1-type formins are associated with actin cables. (A) The evolutionary history of 284 FH2 domain sequences from formin homologs was inferred by the maximum likelihood method for 350 amino acid positions in ≥80% of the sequences. This tree is the same as the tree in **Figure 4**, but includes the FH2 domains from the formins of three additional chytrid species that have been observed to assemble actin cables [*Neocallimastix patriciarum (Li and Heath 1994)*, *Orpinomyces joyonii (Li and Heath 1994)*, and *Chytriomyces hyalinus (Dee et al. 2019)*]. Consensus tree shown, pruned to highlight clade 2 (the main fungal clade) from **Figure 4**. Nodes with <70% bootstrap support were collapsed to polytomies. All bootstrap values are shown. (B) Representative examples of *Spizellomyces punctatus (Sp)* sporangia seeded 1-day prior and *Batrachochytrium salamandrivorans (Bs)* sporangia seeded 3-days prior to being fixed and stained for polymerized actin (alone inverted, black; overlay, green) and for DNA with DAPI (blue). *Sp* sporangia have actin patches (Pa), transient perinuclear actin shells (Ps), and actin cables (Ca). *Bs* sporangia have actin patches and perinuclear actin shells, but no actin cables. (C) Distribution of actin cables and *bni1*-type formins across fungi. Color-filled dots indicate the presence of the given component in the given species. Symbols on the tree represent: fungi (square); Dikarya (pentagon); chytridiomycota (circle). ? indicates the presence of cables in this species is unknown in the literature. * indicates a finding from this paper. *Neo/Neosp1, Neocallimastix patriciarum; Orp/Orpsp1, Orpinomyces joyonii; Bd/BDEND, Batrachochytrium dendrobatidis; Bs/BSALA, Batrachochytrium salamandrivorans; Sp/SPUNC, Spizellomyces punctatus; Rg/RGLOB, Rhizoclosmatium globosum; Ch/Chyhya1, Chytriomyces hyalinus; Am/AMACR, Allomyces macrogynus; Pb/PBLAK, Phycomyces blakesleeanus; Um/UMAYD, Ustilago maydis; Cn/CNEOF, Cryptococcus neoformans; An/ENIDU, Aspergillus nidulans; Sc/SCERE, Saccharomyces cerevisiae; Spo/SPOMB, Schizosaccharomyces pombe*.

### Actin structures in Bd are dynamic and use distinct nucleators

Because actin is required for a wide variety of essential eukaryotic cell functions, we used pharmacological inhibitors to probe actin assembly and dynamics in *Bd* zoospores and sporangia. To determine which actin structures in *Bd* zoospores and sporangia are dynamic, we treated *Bd* for short time periods with Latrunculin B (LatB), a specific small molecule inhibitor of actin polymerization that does not disrupt existing, stable filaments or networks. LatB treatment of zoospores revealed that the four actin phenotypes present in this life stage are dynamic, as the percent of cells displaying pseudopods, spikes, cortical actin, or patches decreased to nearly 0% with 30 minutes of treatment with LatB, compared to the ethanol carrier control (**Fig. 6**).

**Figure 6.**
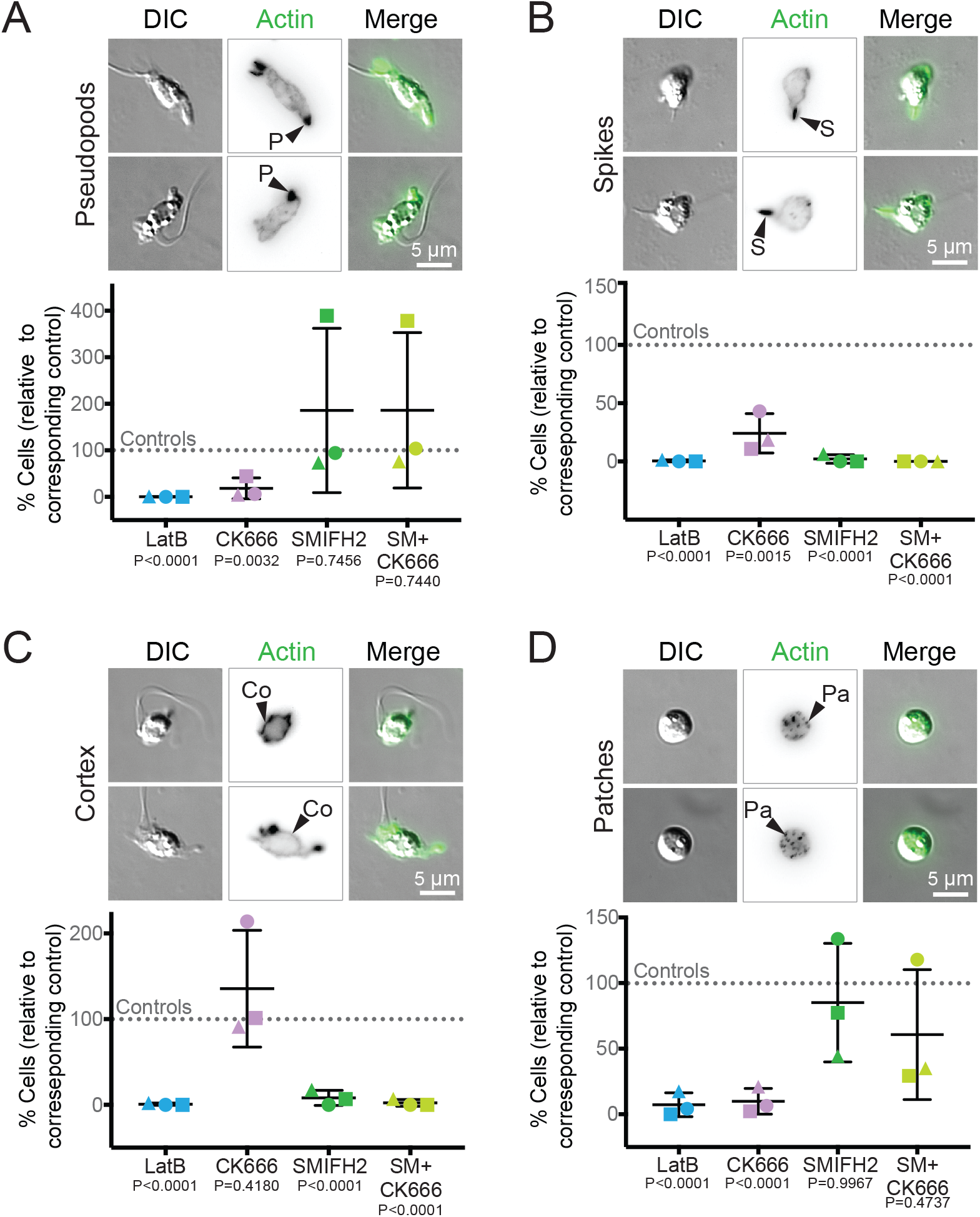
Actin networks in Bd zoospores are dynamic and use distinct nucleators. Synchronized populations of *Bd* zoospores were treated with LatrunculinB (LatB) to identify dynamic actin structures, or with Arp2/3 and/or formin inhibitors for 30 minutes. Cells were then fixed and stained for polymerized actin with fluorescent phalloidin, imaged, and quantified for presence of actin-filled pseudopods (P, part A), actin spikes (S, part B), cortical actin (Co, part C), and actin patches (Pa, part D). Each panel shows examples of cells (DIC: grey) and phalloidin-stained actin structures (alone inverted, black; overlay, green) with relative percent of cells with each structure quantified below. Percent of cells with the indicated actin phenotype for each drug treatment is normalized to its respective control: 1 μM LatrunculinB (LatB) normalized to the ethanol carrier control (EtOH); 100 μM CK666 normalized to its inactive analog CK689; 25 μM SMIFH2 normalized to a DMSO carrier control; and the combination treatment of 25 μM SMIFH2 + 100 μM CK666 (SM+CK666) was also normalized to the DMSO control. For raw data, see **Supplemental File 6**. Three independent experiments were performed, each represented by a different shape, the means and standard deviations shown in black. P-values for each treatment, relative to its respective control, are shown (unpaired Student’s T-tests for EtOH vs. LatB and CK689 vs. CK666; one-way ANOVA with Tukey’s multiple comparisons test for DMSO vs. SMIFH2 and the double treatment). Fluorescent images for (A),(B), and (D) are maximum intensity projections, fluorescent images for (C) are single z-slices to highlight the cortex. Brightness and contrast is not the same across images. Scale bars, 5 μm.

To determine which nucleators are required for building these dynamic actin structures, we then treated zoospores and sporangia with CK666 that inhibits the Arp2/3 complex (Nolen et al. 2009; Hetrick et al. 2013), SMIFH2 that inhibits FH2 domains of formins (Rizvi et al. 2009), or a combination of both. Chytrid Arp2/3 complexes (**Fig. S4**), as well as formin FH2 domains (**Data S3**) are similar to those from animal and fungal systems where these drugs have been previously validated (Nolen et al. 2009; Rizvi et al. 2009; Burke et al. 2014). SMIFH2 was recently shown to also affect some myosins (Sellers et al. 2020), so results from this drug could be due to either formin or myosin inhibition. For each treatment, we blindly measured the frequency of each actin structure in treated samples, and then measured the percentage of cells with each structure. Because of the variability in the prevalence of actin structures between biological replicates (**Fig. 1c**, **Fig. S2**, **Supplemental Data 6**), as well as a slight effect of the drug carrier DMSO on actin phenotype frequencies (**Fig. S2**) we normalized the data from each experimental sample to its control (**Fig 6**, with raw data in **Supplemental Data 6**): 1 μM LatrunculinB (LatB) normalized to the ethanol carrier control (EtOH), 100 μM CK666 normalized to its inactive analog CK689, 25 μM SMIFH2 normalized to a DMSO carrier control, and the combination treatment of 25 μM SMIFH2 + 100 μM CK666 (SM+CK666) was also normalized to the DMSO control.

We first examined the role of Arp2/3 and formins in the assembly of zoospore actin structures. The percent of *Bd* zoospores with pseudopods decreased by an average of 80% with treatment of CK666 (**Fig. 6a**). SMIFH2 had a variable effect on pseudopods; in some trials, the drug had no impact, and in others it drastically increased the percent of cells with pseudopods (**Fig. 6a**). The double treatment showed a similar pattern (**Fig. 6a**). Although the protrusions in SMIFH2 treated zoospores fit our definition of a *Bd* pseudopod (actin-rich and at least 1 μm in width), the protrusions of the SMIFH2-treated cells appear rounder and less protrusive (**Fig. S6**). In contrast, all treatments decreased the percent of cells with actin spikes (**Fig. 6b**). This effect, however, was less drastic with CK666 treatment alone, which decreased spikes by about 70% compared to about 98% for SMIFH2 and 100% for the double treatment (**Fig. 6b**). While the effect of CK666 on the percentage of cells with cortical actin was variable (**Fig. 6c**), the percent of cells with cortex consistently decreased by about 90% with SMIFH2 treatment and about 98% for the double treatment (**Fig. 6c**). We defined a cell with actin patches as one with ≥10 circular localizations of actin; treatment with CK666 reduced the percent of cells with actin patches by an average of 90% (**Fig. 6d**). Treatment with SMIFH2 or the double treatment had large variability in the effect on patch percentage (**Fig. 6d**).

Next we turned to sporangia and found that the perinuclear actin shells and actin patches of *Bd* sporangia have differing stability. Like those in zoospores, the actin patches in sporangia are highly dynamic and nearly disappear upon LatB treatment (8.3±6.3 versus 49.7±23.5 patches/cell in controls, **Figs. 7** and **S7**). Actin patches in sporangia also appear to be Arp2/3-dependent as the average number of patches per cell decreased with CK666 treatment to 22.6±16.43 patches/cell, compared to 58±15.43 patches/cell in CK689 treated control cells (**Fig. 7B**), an effect that was more pronounced in rhizoids (**Fig. 7C**). Because CK666 treatment did not fully match treatment with LatB, we also measured the effects of the formin inhibitor SMIFH2 on actin patch number in sporangia (**Fig. 7**). Treatment with SMIFH2 also reduced the average actin patch number per cell (32.3±13.22), as did treatment with both SMIFH2 and CK666 (20±7.09 patches/cell) compared to the DMSO control (51.3±15.88 patches/cell). Interestingly, the location of the patches in the SMIFH2 treated cells remained relatively unchanged (**Fig. 7A**), though for the double treatment, patches were severely reduced in the rhizoids (**Fig. 7C**). Perinuclear shells appear only partially dynamic, as a portion of the shell remains after 30 minutes of LatB treatment (**Fig. 7**, **Fig S7**).

**Figure 7.**
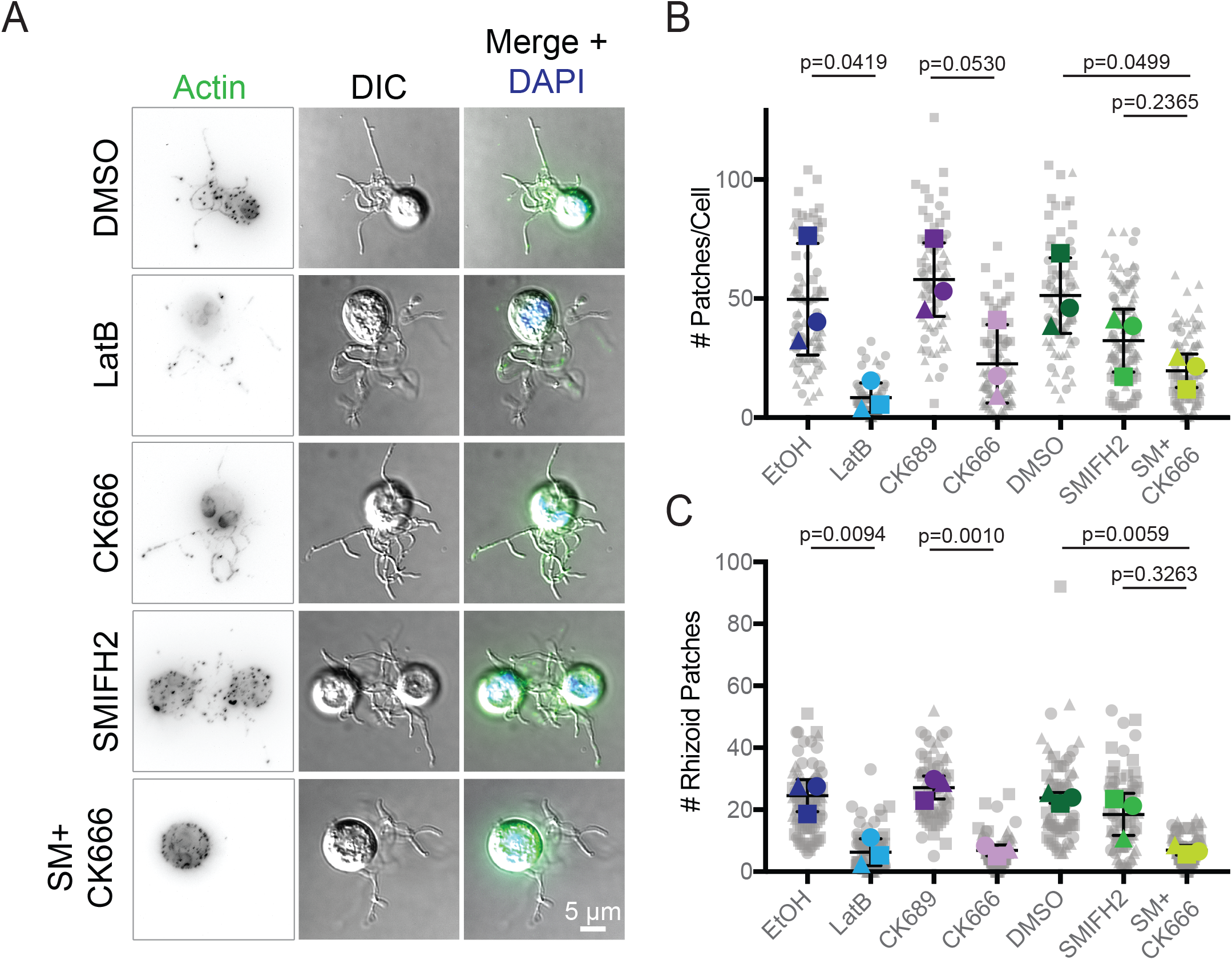
Actin patches in Bd sporangia are dynamic and use the Arp2/3 complex. Populations of *Bd* sporangia seeded 1 day prior were treated with drugs using the same concentrations as in **Figure 6**, then fixed and stained for polymerized actin with phalloidin and for DNA with DAPI. (A) Examples of sporangia (DIC: grey) and phalloidin stained actin patches (inverted, black; green in overlay), with an overlay including the nucleus (blue) after treatment with each drug. Though DMSO has a slight effect on patches (see **Fig. S2**), all controls looked phenotypically similar (see **Fig S7**), so only a DMSO treated cell is shown. (B) quantification of the number of actin patches per sporangia. Larger, colored shapes indicate the average number of patches per cell in each treatment from three independent experiments, each represented by a different shape. Each gray shape represents the number of patches in a single cell in that experiment. Means and standard deviations of these averages are shown in black. Statistical tests were performed using the averages of the three experiments (i.e., the three colored shapes). P-values for each treatment, relative to its respective control, are shown (unpaired Student’s T-tests for EtOH vs. LatB and CK689 vs. CK666; one-way ANOVA with Tukey’s multiple comparisons test for DMSO vs. SMIFH2 and the double treatment). Brightness and contrast is not the same across images. Scale bar, 5 μm.

These results show that most actin structures in *Bd* are dynamic and suggest that the Arp2/3 complex contributes to the formation of pseudopods and patches, while formins are used to build the cortex, and both appear to help build actin spikes.

## DISCUSSION

Chytrids share a number of traits with other opisthokonts that are not found in the Dikarya, including microtubule-based flagella (Longcore et al. 1999; Hodges et al. 2010), cells that lack cell walls, and both animal-typical and fungal-typical cell cycle control machinery (Medina et al. 2016). Here, we show that chytrid fungi have an actin regulatory protein repertoire that also appears intermediate to that of animals and Dikarya. For example, while animals, chytrids, and Dikarya all have a complete Arp2/3 complex, some of its activators and other actin regulatory proteins are conserved in animals and chytrids but are missing in the Dikarya. This includes all members of the SCAR/WAVE complex, DAAM formins, gelsolin/villin family proteins, talin, and Myo22 (**Fig. 2**, **Fig. S4**). There are also a number of proteins which have been lost throughout the fungal lineage, such as the WASH complex and ENA/VASP family proteins (**Fig. 2**, **Fig. S4**). Many of these regulators are thought to have been present in the last common eukaryotic ancestor, indicating that they were lost during fungal evolution (Veltman and Insall 2010; Kollmar et al. 2012). Our finding of key actin associated proteins in chytrid fungi that have been lost in Dikarya highlights the potential for using chytrid fungi to explore actin cytoskeletal evolution.

Our analysis of formin evolution largely recapitulates the topologies reported in previous phylogenies (Higgs and Peterson 2005; Chalkia et al. 2008; Pruyne 2017), but with greater taxonomic diversity and support. While the formins in some well-supported clades have similar domain architectures (**Fig. 4**, clade 1 has diaphanous-like architectures), other clades unite proteins whose domain structures vary wildly (**Fig. 4**, clade 7 has diaphanous-like and inverted-like architectures). The scattering of domain architectures throughout the phylogeny indicates that domain shuffling is common in the formins, particularly with PTEN/PTEN-like formins, which arose from at least two independent events (Pruyne 2017; van Gisbergen et al. 2018).

Our formin phylogeny shows that, like other actin regulators, chytrids share formin families found in animals that are missing from Dikarya, including DAAM and Delphilin formins (**Fig. 4**). Inclusion of multiple chytrid species in our analysis also revealed variability in formin content among chytrid species. This variation includes a remarkable correlation between the presence of *bni1*-type formins and fungal cells that build actin cables (**Fig. 5**). This suggests that *bni1*-type formins may be required specifically to build actin cables. *Bd*, a species that lacks *bni1*-type formins, still builds spikes, pseudopods, cortical actin, and other actin structures, indicating that their assembly does not require *bni1-*type formins. The presence of ancestral formins alongside *bni1-*type formins would further suggest that cables evolved within organisms capable of building animal-like actin networks and were retained in Dikarya while the other formins were lost.

*Bd* zoospores assemble a variety of dynamic actin structures whose appearance and regulation appear similar to those of human cells. In addition to Arp2/3-dependent pseudopods (**Fig. 6a** and (Fritz-Laylin, Lord, et al. 2017), *Bd* zoospores build thin, actin-filled protrusions we call spikes. Like the thin, actin-filled filopodia of animal cells that require both Arp2/3 and formin activity (Svitkina et al. 2003; Pellegrin and Mellor 2005; Schirenbeck et al. 2005; Harris et al. 2010; Block et al. 2012), spikes in *Bd* are sensitive to both CK666 and SMIFH2 (**Fig. 6**). Although shorter than animal (on average 5-20 μm, (Gallop 2020)) and *Dictyostelium* filopodia (2-5 μm, (Medalia et al. 2007)), the visual similarity as well as the apparent involvement of both Arp2/3 complex and formin protein activity, raises the possibility that these actin spikes are related to filopodia. In the future, we will look for evidence of actin monomer addition to the tips of these structures coupled with outward protrusion—a key behavior of filopodia. *Bd* zoospores also assemble an actin cortex that is sensitive to SMIFH2 treatment (**Fig. S1**). Because the formin inhibitor SMIFH2 has recently been shown to also inhibit some myosins (Sellers et al. 2020), it is possible that the reduction in spike formation and cortex intensity upon SMIFH2 treatment are due to inhibiting formins, myosins, or a combination of both. Taken together, our results indicate that *Bd* zoospores assemble actin structures similar to those of human cells.

While the zoospore actin cytoskeleton resembles that of human cells, the actin cytoskeleton of *Bd* sporangia more closely resembles that of yeast and other Dikarya, particularly in the assembly of small actin patches near the cell periphery (**Fig. 1d**). Based on their similarity to actin patches of yeast and other Dikarya, we predict that these actin patches are involved in endocytosis and cell wall synthesis, explaining their abundance in growing sporangia. We hypothesize that the patches seen in a minority of zoospores represent cells that have initiated their transition to the sporangial growth stage (**Fig. 1b**). Like actin patches of yeast, the actin patches of *Bd* sporangia are sensitive to Arp2/3 inhibition (**Fig 7b**). Actin patches may also require formins, but the effect of SMIFH2 is variable (**Fig. 6d**, **Fig. 7b**). *Bd* sporangia also build perinuclear actin shells (**Fig. 1d**), similar to those found in other species of chytrid fungi. Earlier reports of perinuclear shells suggested they were fixation artifacts (Li and Heath 1994), but recent live imaging of actin in *Sp* clearly shows that these structures are present in living sporangia and form just before mitosis (Medina et al. 2020).

We know of no Dikarya cells with the animal-like actin phenotypes seen in chytrid zoospores. Chytrids use these structures for crawling, a behavior not seen in sessile fungi that spend their life cycle enclosed in cell walls. The simplicity of the actin cytoskeleton in Dikarya, therefore, may have corresponded with the loss of the zoospore stage type that uses these actin regulators to build the animal-like actin structures.

Studying chytrid actin networks provides important clues about the evolution of Dikarya and also gives us valuable information about chytrid cell biology. *Bd* is a causative agent of Chytridiomycosis, a deadly skin infection of amphibians that is associated with population declines around the world (Berger et al. 1998; Longcore et al. 1999; Lips 2016; O’Hanlon et al. 2018). Although the exact mechanism of infection by *Bd* is unknown, the main model is that zoospores attach to the host, encyst, and the sporangia invade deeper skin layers as they grow (Van Rooij et al. 2015). The sporangia mature as the frog sheds its skin, allowing the release of zoospores back into the environment (Van Rooij et al. 2015). We confirmed our previous finding of pseudopods and cortical actin in *Bd* (Fritz-Laylin, Lord, et al. 2017), and discovered new actin structures not seen in *Bd* before: spikes and patches. These structures could play important roles in the infection process. For example, zoospores could use pseudopods to crawl along the surface of the host to find a suitable local environment before encysting. Spikes, if they function like filopodia, could also be used for movement or for sensing local environmental conditions. Additionally, actin patches likely represent the endocytic sites necessary for the massive nutrient uptake needed for rapid growth of sporangia and production of new zoospores. This model suggests that actin networks underlie the motility and rapid growth that are key to the pathology and pathogenicity of *Bd*.

## MATERIALS AND METHODS

### Identification of actin regulatory proteins

Protein sequences from the following chytrids were retrieved from the website of the National Center for Biotechnology Information (NCBI; ncbi.nlm.nih.gov/): *Batrachochytrium dendrobatidis* (*Bd*) strains JAM81 (GCA_000203795.1, JGI-PGF project ID 4001669; http://genome.jgi-psf.org/Batde5) and JEL423 ((Farrer et al. 2017), *Batrachochytrium salamandrivorans* (*Bs;* GCA_002006685.1, (Farrer et al. 2017)*; Spizellomyces punctatus DAOM BR117* (*Sp; GCA_000182565.2*, (Russ et al. 2016)*; Rhizoclosmatium globosum JEL800* (*Rg; GCA_002104985.1*, (Mondo et al. 2017); and *Allomyces macrogynus ATCC 38327* (*Am;* GCA_000151295.1, Broad Institute). Homologs of proteins from the model organisms used in this study [*Saccharomyces cerevisiae S288C (Sc), Schizosaccharomyces pombe* strain 972 (*Spom*) *Dictyostelium discoideum AX2 and AX4 (Dd)*, *Homo sapiens, Mus musculus (*occasionally used to confirm results), and *Arabidopsis thaliana (At)*] were identified using a combination of the Basic Local Alignment Search Tool (BLAST; (Altschul et al. 1990)), literature review, and probing Swiss-Prot reviewed entries on UniProtKB. Individual UniProtKB IDs for proteins from these model species are provided in **Table S1**, along with the NCBI accession numbers for the indicated chytrid species. Additional sequences were retrieved from NCBI from species used for actin identification [*Oryctolagus cuniculus*, *Giardia intestinalis (GIAIN), Chlamydomonas reinhardtii (CHLRE)].* Multiple splice variants were not included in this analysis.

#### Actin

BLASTp using standard parameters (E= 1.0×10^−5^, word size=3, BLOSUM 62 matrix, filtering low complexity regions) and Rabbit, *Sc, Spom*, and *Dd* actin sequences as queries was used to identify actin homologs in the five chytrid species of interest. Chytrid sequences sharing ≥50% sequence identity with any of these queries were compiled and aligned with the query sequences as well as actin sequences from *GIAIN* and *CHLRE*; Arp1 from human, *Sc*, *Spom*, and *Dd*; and Arp2 and Arp3 from human, *Sc*, *Spom*, *Dd*, and the five chytrids. With this alignment, a simple Maximum Likelihood phylogeny was built using the IQtree web server (default parameters; (Trifinopoulos et al. 2016)). Chytrid actin sequences were defined as those which formed a clade with known actin sequences.

#### Actin regulatory proteins

BLASTp with standard parameters (E= 1.0×10^−5^, word size=3, BLOSUM 62 matrix, and filtering low complexity regions) and queries from *Sc*, *Spom*, *Dd*, mouse, or human was used to identify homologs for 34 actin regulatory proteins, complexes, or protein families in the five chytrid species of interest. BLAST hits were confirmed by obtaining a mutual best BLAST hit (MBBH) and identifying the predicted domain structure of all potential chytrid homologs. MBBHs were found through NCBI (same parameters) or yeastgenome.org (TBLASTN, default parameters, open reading frames dataset; (Cherry et al. 2012) using the potential chytrid homologs as search queries. The predicted domains of all potential chytrid homolog sequences were obtained using the hmmscan tool (hmmer.org) against the Pfam database (v32; (El-Gebali et al. 2019) using the website of the European Bioinformatics Institute (EMBL-EBI; https://www.ebi.ac.uk/) or on the command line using hmmscan from the hmmer suite v3.2.1 (hmmer.org) and the PfamA Hidden Markov Model (v32; (El-Gebali et al. 2019). In some cases, MBBHs were not obtained due to the complexity of the protein family, but domain architectures and multiple sequence alignments of the chytrid homologs confirm their family membership.

#### Myosin

Full length myosin protein sequences for *Allomyces macrogynus* (ATCC 38327), *Batrachochytrium dendrobatidis* (JAM81), *Spizellomyces punctatus* (DAOM BR117) *Saccharomyces cerevisiae* (RM11-1a), *Schizosaccharomyces pombe* (972h), *Arabidopsis thaliana, Dictyostelium discoideum* (AX4) and *Homo sapiens* were obtained from Cymobase (www.cymobase.org) (Odronitz and Kollmar 2006). Myosin sequences for *Batrachochytrium salamandrivorans* and *Rhizoclosmatium globosum* were identified by extensive BLAST searches using either full-length, myosin motor domain or MyTHFERM domain from Bd myosins as a query. The classification of the myosins was validated by a general BLAST search of the full protein and manual inspection of the tail domains.

### Identification of FH2 domain-containing proteins

Identifying potential formin sequences required a different approach from the rest of the actin regulatory proteins. The predicted FH2 domains from *Sc* formins Bni1p and Bnr1p were isolated and used as independent search queries using Position Specific Iteration-BLAST (PSI-BLAST; (Altschul et al. 1997) through NCBI (default parameters, filter low complexity regions). For each chytrid species and for each search query, five iterations of PSI-BLAST were run, after which convergence occurred and subsequent iterations yielded no new sequences above the E-value threshold (0.005). All sequences above the threshold after these iterations were checked for an FH2 domain using the domain prediction method from above; those without an FH2 domain were removed from the dataset. Surrounding gene annotations of proteins with only FH2 domains were checked to identify other formin-typical domains to create a full length formin sequence. Splice variants were not included in this analysis.

### Identification of formin domain organizations

All known formin sequences from human, *Sc, Spom, Dd*, and *A*t were checked for predicted domains using the same domain prediction method as above and confirming these results with the literature. All chytrid FH2-domain containing sequences were checked for predicted domains as well. Coiled coil regions, signal peptides (SigPs) in plant formins, and transmembrane (TMs) in plant formins were determined using this method, as EBI runs hmmscan, a coiled coil prediction algorithm (Lupas et al. 1991), and the Phobius program (Käll et al. 2004) for SigPs and TMs simultaneously. FH1 domains contain many polyproline stretches and are hard to determine through computational methods, thus FH1 domains in chytrid sequences were determined by hand. Polyproline stretches were defined as being at least 6 prolines long out of 7 consecutive residues, the minimum number of prolines needed to bind profilin (Perelroizen et al. 1994; Petrella et al. 1996; Mahoney et al. 1999; Paul and Pollard 2008). An FH1 domain was defined as the stretch of amino acids from the first proline of the first polyproline stretch to the last proline of the last polyproline stretch directly N-terminal to the FH2 domain. The diaphanous autoregulatory domain of diaphanous like formins is short and also needed to be identified by hand. This domain is C-terminal to the FH2 domain and has the consensus sequence MDXLLXXL (Alberts 2001; Higgs 2005). The C-termini of all chytrid formins were aligned using TCoffee (default parameters) and checked for the consensus sequence. Sequences with obvious insertions or deletions in the diaphanous autoregulatory domain region were removed and the remaining sequences were realigned and reexamined.

### Phylogenetic analysis of formin proteins

#### Main analysis

FH2-domain-containing sequences were obtained from the species distribution sunburst of the FH2 domain page (PF02181) on the Pfam website (v32; pfam.xfam.org; (El-Gebali et al. 2019). The following species in the given taxa were used in this analysis: <u>Chytridiomycota</u>: *Bd, Bs, Sp, Rg;* <u>Blastocladiomycota</u>: *Am*; <u>Mucoromycota</u>: *Phycomyces blakesleeanus*; <u>Dikarya</u>: *Sc, Spom*, A*spergillus nidulans, Cryptococcus neoformans*, U*stilago maydis*; <u>Microsporidia</u>: *Spraguea lophii*; <u>Metazoa</u>: Human, mouse, *Drosophila melanogaster*; <u>Choanoflagellates</u>: *Monosiga brevicollis, Salpingoeca rosetta*; <u>Filasterea</u>: *Capsaspora owczarzaki*; <u>Apusozoa</u>: *Thecamonas trahens*; <u>Amoebozoa</u>: *Dd*, *Acanthamoeba castellanii, Enantoboeba invadens*; <u>Discoba</u>: *Naegleria gruberi, Trypanosoma brucei*; <u>SAR</u>: *Thalassiosira pseudonana, Plasmodium falciparum,Tetrahymena thermophila*; <u>Plants</u>: *At, CHLRE, Physcomitrella patens*; <u>Metamonads</u>: *Trichomonas vaginalis.* The CD-hit program (Huang et al. 2010; Fu et al. 2012) was used to remove sequences which were ≥98% identical, reducing redundancy in the data set. The remaining sequences were aligned to the full HMM for the FH2 domain from Pfam using hmmalign (no additional program options) from the hmmer suite v3.2.1. The FH2 domains were isolated by trimming the alignment according to the FH2 domain of the S*c* Bni1p sequence (starting PHKKLKQ; ending ADFINEY), which had been previously hand clipped and provided an accurate judgement for the placement of other FH2 domains. Columns which had <80% occupancy were removed from the alignment. A Maximum Likelihood phylogeny was generated from this curated alignment using the IQTree webserver (default parameters; (Trifinopoulos et al. 2016).

#### bni1-type formin analysis

We searched the literature and identified the following chytrid species with observed actin cables: *Neocallimastix patriciarum (Li and Heath 1994)*; *Orpinomyces joyonii (Li and Heath 1994)*; and *Chytriomyces hyalinus (Dee et al. 2019)*. *N. particiarum* and *O. joyonii* did not have genomes in the JGI database, so we used the genomes of species in the same genus, assuming that the presence of cables is consistent across a genus. We used the FH2 domains from *Bd* JAM81 formins as queries for TBLASTN searches (default settings, perform gapped alignments) against the filtered model transcripts database for the following species’ genomes on JGI MycoCosm: *Neocallimastix californiae* G1 (Haitjema et al. 2017); *Orpinomyces sp.* strain C1A v1.0 (Youssef et al. 2013); Chytriomyces hyalinus JEL632 v1.0 (JGI, https://mycocosm.jgi.doe.gov/Chyhya1/). All hits were confirmed for the presence of an FH2 domain using the same method described above. FH2-domain-containing sequences from these three species were added to the file containing the formin sequences used in the main analysis and the same alignment, clipping, column editing, and tree building processes were performed as before. The large tree was pruned using the iTOL website (v5.5.1; (Letunic and Bork 2019) to show only the main fungal clade that included Bni1p and Bnr1p from *Sc*.

### Cell culture and population synchronization

Cultures of *Bd JEL423* were maintained in 1% Tryptone broth at 24°C in non-ventilated, tissue-culture treated flasks. For experiments investigating zoospores, populations seeded 3 days prior were synchronized to ensure the cells were in a similar developmental stage. Synchronization was achieved by washing out previously-released zoospores 3 times with 1% Tryptone and adding fresh Tryptone for the sporangia left adhered to the flask. The sporangia were incubated at 24°C for 2 hours before newly released zoospores were harvested from suspension via centrifugation at 2000xg for 5 minutes. For experiments investigating sporangia, cells from populations seeded in non-tissue culture treated flasks 1 day prior (called 1-day cultures) were harvested from suspension via centrifugation at 2000xg for 5 minutes.

### Chemical inhibition of actin and actin nucleators

Synchronized zoospores were adhered to the bottom of 96-well plates using 0.5 mg/mL Concanavalin A (Sigma, C2010). Adhered cells were then treated with Bonner’s Salts (Bonner 1947), 1uM LatrunculinB (Millipore, 4280201MG) or an equal volume of ethanol, 100uM CK666 (Calbiochem/Sigma, 182515) or 100uM of the inactive analog CK689 (Calbiochem/Sigma, 182517), 25uM SMIFH2 (Tocris Bioscience, 440110) or equal volumes of DMSO, and 100uM CK666 + 25uM SMIFH2. Cells were fixed using fixation buffer (4% PFA and 50 mM Sodium Cacodylate, pH=7.2) on ice for 20 minutes, permeabilized and stained for DNA using a mixture of 0.1% Triton X and DAPI (0.5 μg/ml; Life Technologies, D1306) in PEM buffer (100mM PIPES, 1mM EGTA, 0.1mM MgSO4) at room temperature for 10 minutes, and then stained for polymerized actin using Alexa Fluor488 Phalloidin (0.2 U/mL; Life Technologies, A12379) at room temperature for at least one hour. Sporangia from 1-day cultures were adhered to the bottom of 96-well plates using ~0.1% Polyethyleneimine (Sigma, P3143) and then treated, fixed, and stained using the same procedure used for the zoospores.

### Microscopy and image analysis

Cells were imaged on an inverted microscope (Ti-2 Eclipse; Nikon) with a 100X 1.45 NA oil objective and using NIS Elements software. Images were taken using both differential interference contrast (DIC) microscopy and widefield fluorescence microscopy with 460 nm to visualize phalloidin and 360 nm light to visualize DAPI. All images were taken in Z-stacks to encompass the whole cell. All imaging was done at room temperature in PEM buffer. Image processing and analysis was performed in Fiji (Schindelin et al. 2012), and blind scoring was performed using the CellCounter plugin (Kurt De Vos, https://imagej.nih.gov/ij/plugins/cell-counter.html). Zoospores were categorized based on the actin structures present in each cell: pseudopods, actin spikes, cortical actin, actin patches, or a combination of any of these. Pseudopods were defined as bright areas of actin staining 1-2 μm wide, while spikes were defined as being less than 1 μm wide and at least 1 μm long. Cortical actin was defined as bright staining for actin along the outer edge of at least 50% of the cell. Actin patches were defined as at least 10 bright spots of actin <1 μm in diameter in the cell. The number of patches in each sporangium was counted and patch position (cell body vs. rhizoid) was noted.

For zoospores, the percent of cells with each actin structure for each treatment was normalized to its respective control: LatrunculinB (LatB) normalized to the ethanol carrier control (EtOH), CK666 normalized to its inactive analog CK689, SMIFH2 normalized to a DMSO carrier control, and the combination treatment SMIFH2 + CK666 (SM+CK666) was also normalized to the DMSO control. Statistical tests were performed on these normalized values and their respective control, for the three independent experiments. For sporangia, the average number of patches per cell in each treatment was calculated for three independent experiments. Statistical tests were performed using the averages of the three experiments. For all statistical tests, we used unpaired Student’s T-tests for EtOH vs. LatB and CK689 vs. CK666 and a one-way ANOVA with Tukey’s multiple comparisons test for DMSO vs. SMIFH2 and the double treatment.

## ACKNOWLEDGEMENTS

We would like to thank Tom Pollard, Harry Higgs, and Jessica Henty-Ridilla for feedback on Figure 2, as well as Madelaine Bartlett, Edgar Medina, Katrina Velle, and Samuel Lord for comments on the manuscript. This material is based upon work supported by the Pew Charitable Trust (to L.K.F.-L.) and the National Institutes of Health (R01GM12291 to M.A.T.).

---

**Figure S1.**
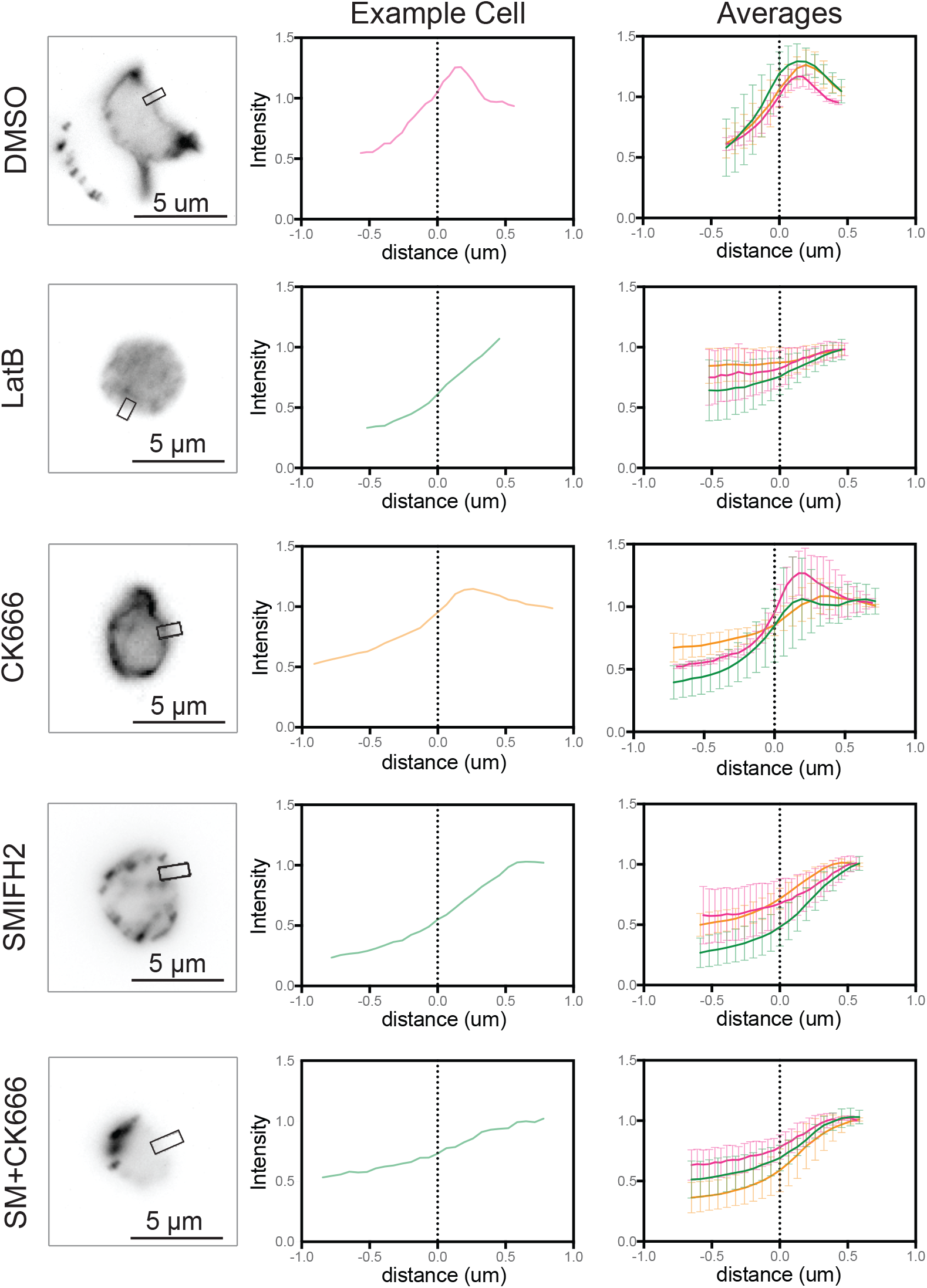
The actin cortex of Bd zoospores is dynamic and is affected by SMIFH2. Synchronized populations of *Bd* zoospores were treated with various actin inhibitors for 30 minutes (See **Fig. 6** for concentrations). Cells were then fixed and stained for polymerized actin with fluorescent phalloidin, imaged, and line scans perpendicular to the cell edge were taken. Actin structures (inverted, black) in representative cells with the location of the line scan overlayed (left). Phalloidin intensity normalized to the cell interior plotted against distance along the line normalized with the cell edge at 0 μm for the cell shown (middle). Average intensities and standard deviations for three independent experiments (N=6 cells), normalized in the same way (right). Scale bars, 5 μm.

**Figure S2.**
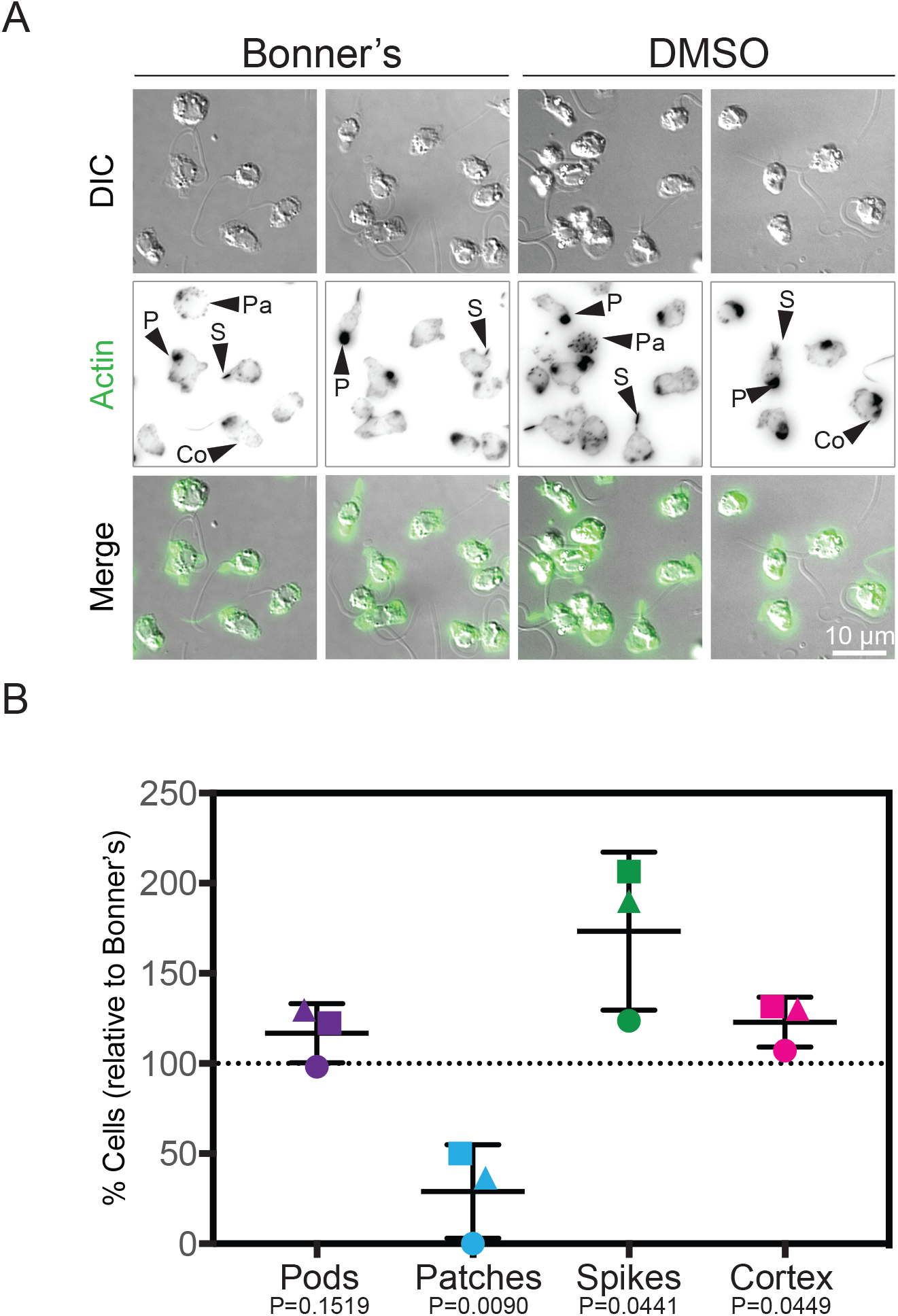
DMSO has a slight effect on the actin structures in Bd. Synchronized populations of *Bd* zoospores were treated with DMSO at the same concentration used in **Figure 6**, or equal volume of Bonner’s salts as a control. Cells were then fixed and stained for polymerized actin with fluorescent phalloidin, imaged, and quantified for presence of actin-filled pseudopods (P), actin spikes (S), cortical actin (Co), and actin patches (P). (A) Representative examples of cells (DIC, grey), and phalloidin stained actin structures (inverted, black), with an overlay of the two (actin, green) after treatment with Bonner’s salts or DMSO. (B) Quantification of the percent of cells with each structure in the DMSO treated cells, normalized to the Bonner’s salts control. P-values for each structure, relative to the Bonner’s control, are shown (unpaired Student’s T-tests). All fluorescent images are not at the same brightness and contrast scale. Scale bar, 10 μm.

**Figure S3.**
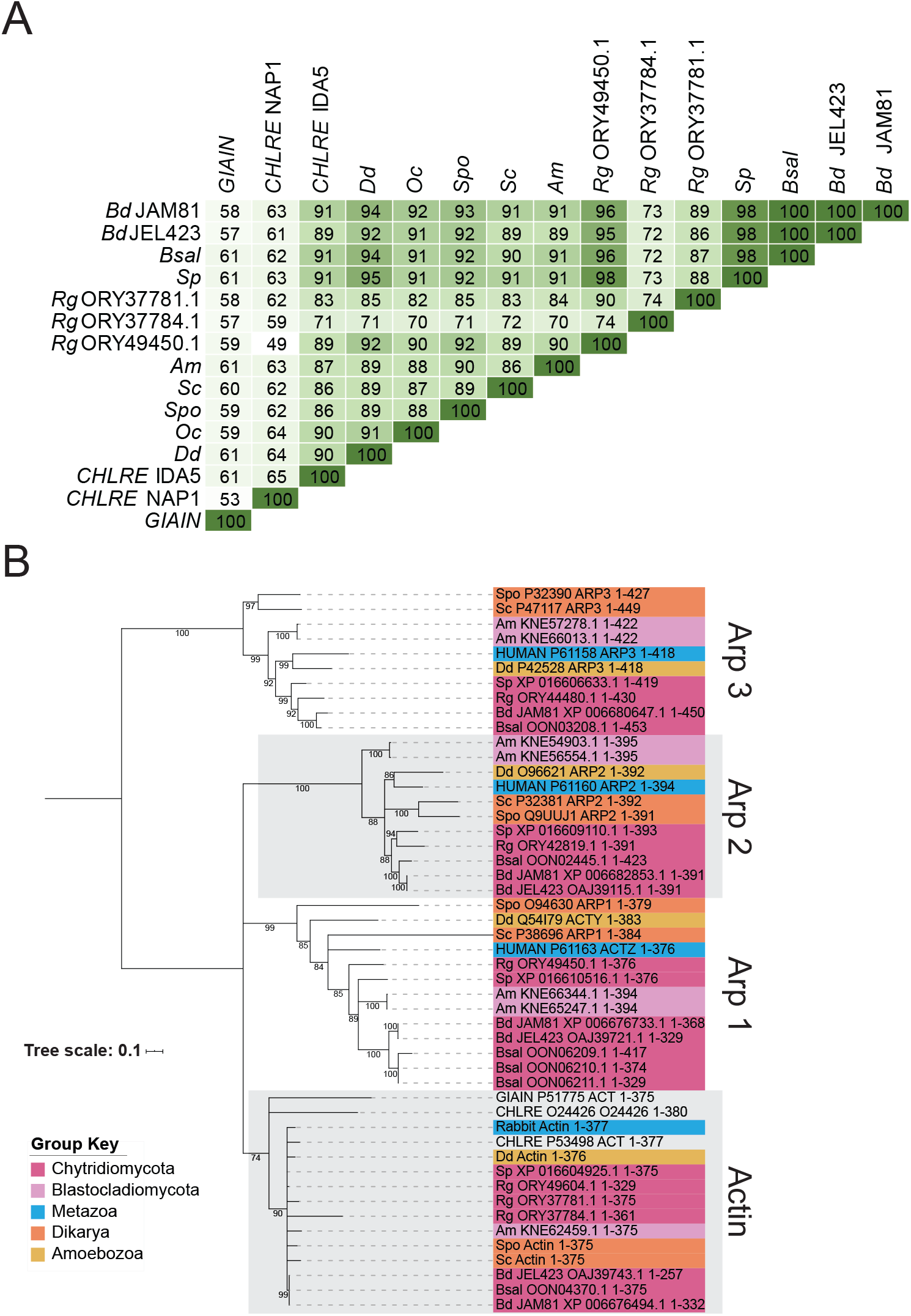
Chytrids have canonical actin. All chytrid sequences sharing ≥50% similarity with Rabbit, *Dictyostelium discoideum*, *Saccharomyces cerevisiae*, or *Schizosaccharomyces pombe* actin sequences were obtained using BLAST and aligned with known actin, Arp1, Arp2, and Arp3 sequences from various taxa. (A) Percent similarities between the actin sequences of the given species. (B) Maximum likelihood phylogeny built to determine which chytrid sequences were true actin sequences. Consensus tree shown, branches with less than 70% bootstrap value were collapsed to polytomies. All bootstrap values shown. True chytrid actin sequences were determined as those that grouped with known actin sequences from other organisms. Rabbit/*Oc*, *Oryctolagus cuniculus*, HUMAN, *Homo sapiens, Dd, Dictyostelium discoideum; Bd, Batrachochytrium dendrobatidis* (JAM81 and JEL423 strains); *Bs, Batrachochytrium salamandrivorans; Sp, Spizellomyces punctatus; Rg, Rhizoclosmatium globosum; Am, Allomyces macrogynus; Sc, Saccharomyces cerevisiae; Spo, Schizosaccharomyces pombe; CHLRE, Chlamydomonas reinhardtii; GIAIN, Giardia intestinalis*.

**Figure S4.**
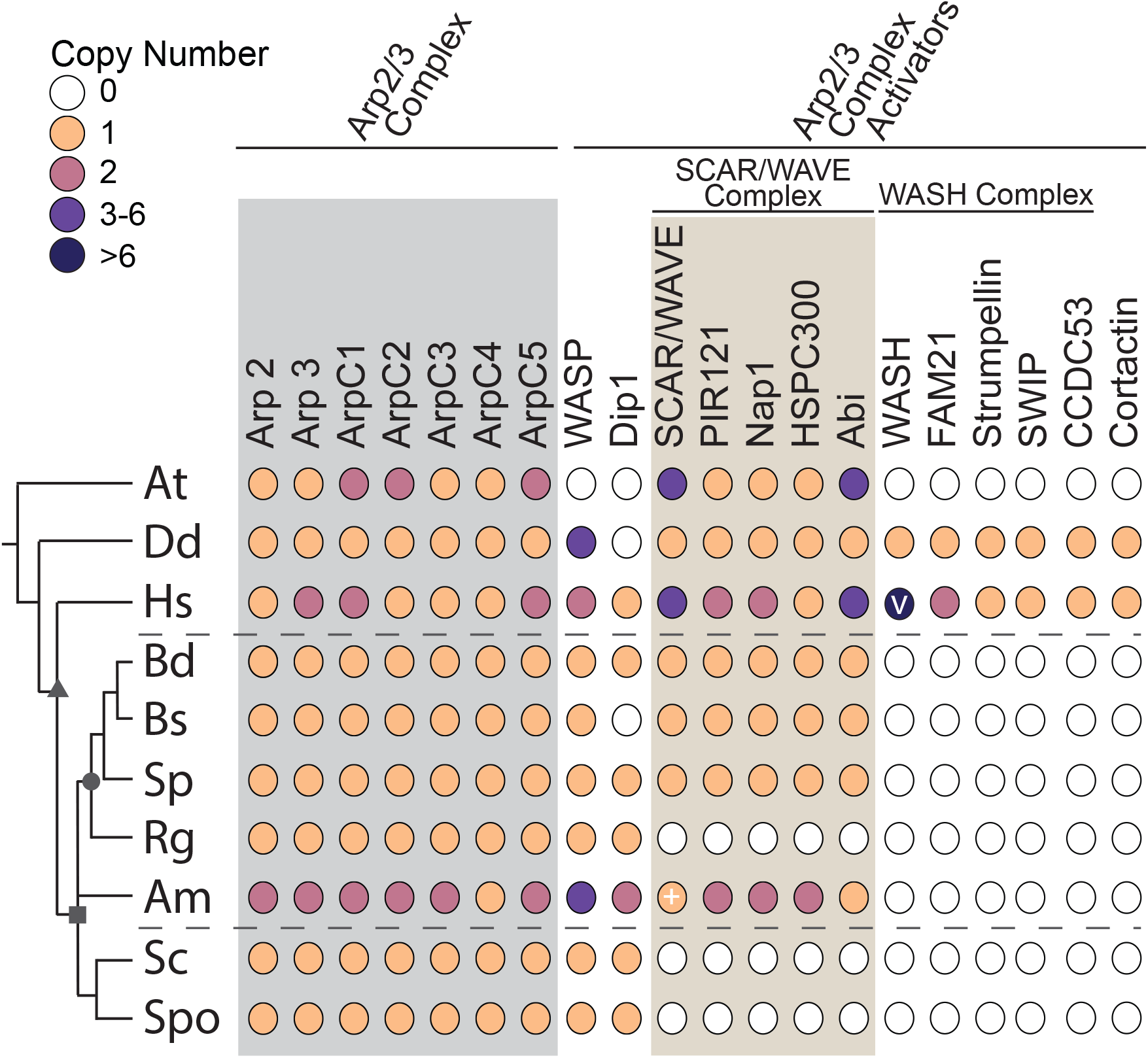
Chytrids retained the Arp2/3 complex and many of its regulators. The distribution of the Arp2/3 complex and its regulators across taxa. Color-filled circles indicate the presence of clear homologs found, with the number of homologs for each protein in each species shown in the colors specified in the key, whereas white dots indicate that no homolog was detected in that species. Dashed lines mark the chytrids. Symbols on the tree represent: opisthokonts (triangle); fungi (square); chytridiomycota (circle). V, copy number of WASH varies individually, as many WASH genes are subtelomeric (Linardopoulou et al. 2007). +, See **Supplemental Table 1** for details and additional potential homologs with caveates. *At, Arabidopsis thaliana; Dd, Dictyostelium discoideum; Hs, Homo sapiens; Bd, Batrachochytrium dendrobatidis; Bs, Batrachochytrium salamandrivorans; Sp, Spizellomyces punctatus; Rg, Rhizoclosmatium globosum; Am, Allomyces macrogynus; Sc, Saccharomyces cerevisiae; Spo, Schizosaccharomyces pombe*.

**Figure S5.**
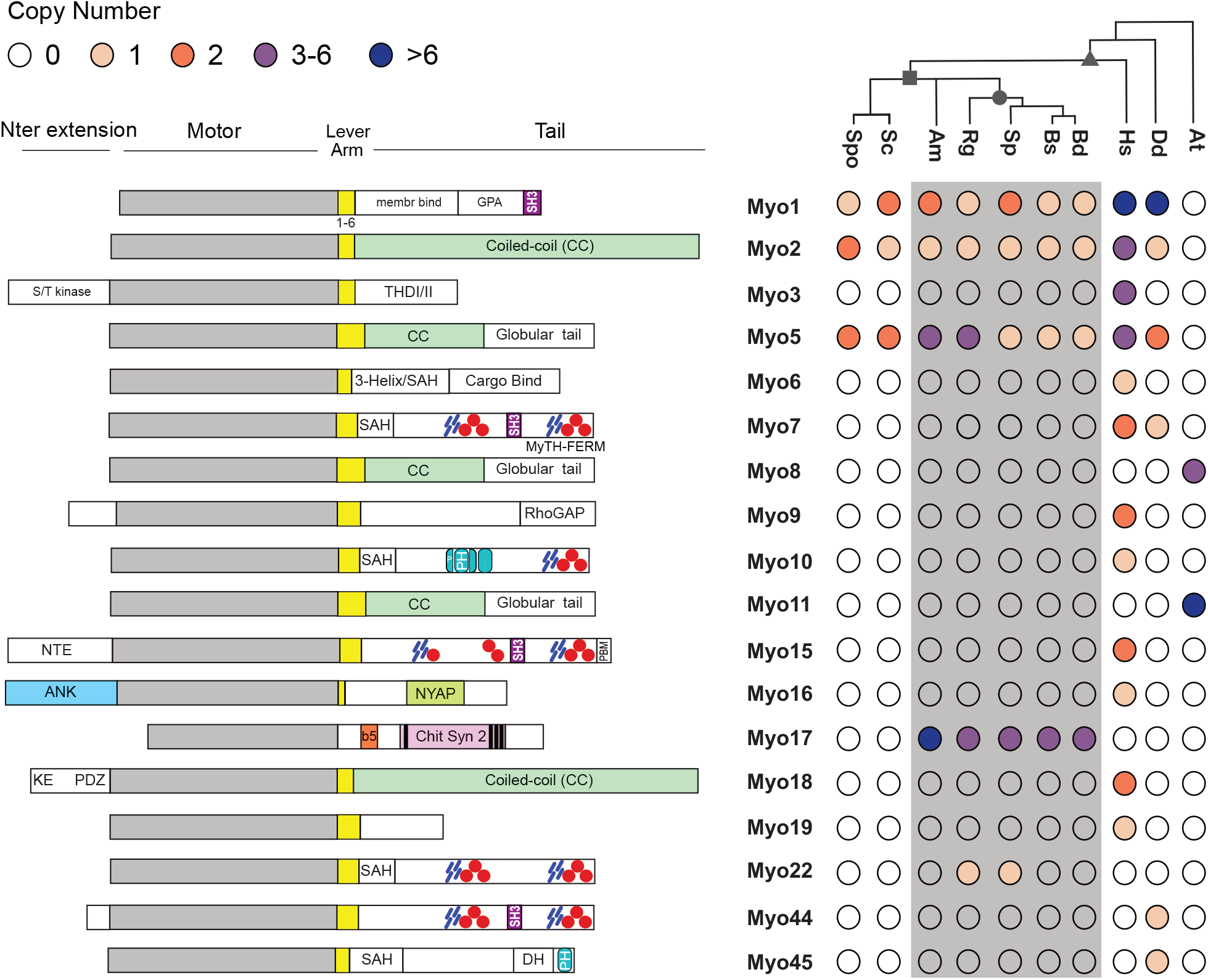
Chytrids have myosins from at least 4 classes. The distribution of given myosins (left, not to scale), across taxa (right). Color-filled circles indicate the presence of at least one myosin of that class in that species, with color indicating the number of myosins. Unfilled circles indicate that no myosin of that class was found in the given species. The domain abbreviations are as follows: yellow box, light chain binding domains (IQ motifs) that can vary in number; blue rectangles, MyTH4 domain; red circles, FERM domain; light blue oval, PH domain; black bars, transmembrane domain; membr binding, membrane binding; GPA, glycine-proline-alanine rich; SH3, src homology 3 (purple rectangle); S/T kinase, serine-threonine kinase; THDI/II, tail homology domain I/II; 3-Helix, triple helix bundle; SAH, single alpha-helix; NTE, N-terminal extension; PBM, PDZ binding motif; ANK, ankyrin repeats; NYAP, Neuronal tYrosine-phosphorylated Adaptor for the PI 3-kinase domain; b5, cytochrome heme binding domain; Chit syn 2, chitin synthase 2; KE, lysine and glutamate rich; PDZ, PDZ domain; DH, Dbl homology domain. *At, Arabidopsis thaliana; Dd, Dictyostelium discoideum; Hs, Homo sapiens* (non-muscle myosins only depicted)*; Bd, Batrachochytrium dendrobatidis; Bs, Batrachochytrium salamandrivorans; Sp, Spizellomyces punctatus; Rg, Rhizoclosmatium globosum; Am, Allomyces macrogynus; Sc, Saccharomyces cerevisiae; Spo, Schizosaccharomyces pombe*.

**Figure S6.**
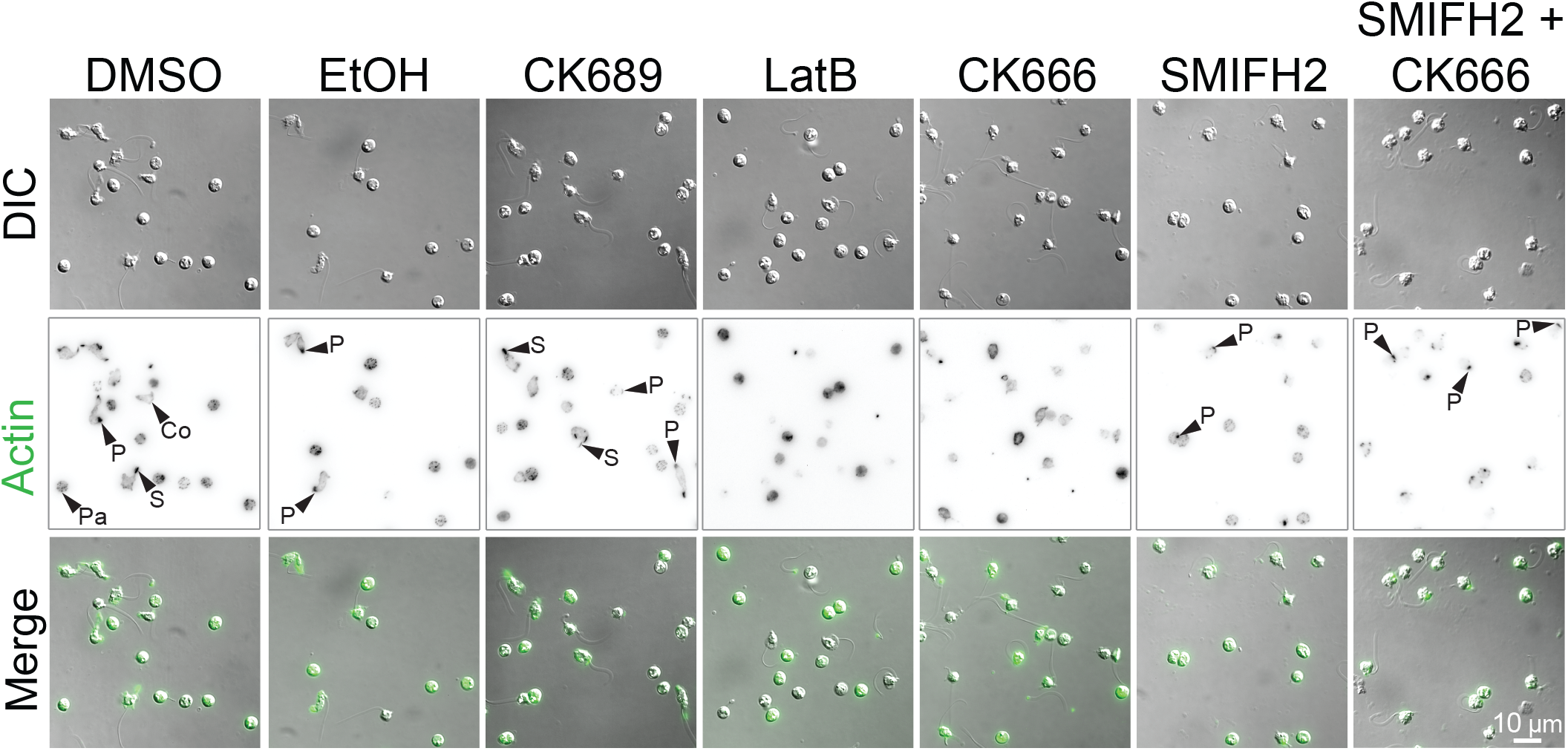
Additional examples of actin structures in Bd zoospores. Synchronized populations of *Bd* zoospores were treated with various actin inhibitors for 30 minutes (see **Figure 6** legend for concentrations). Cells were then fixed and stained for polymerized actin with fluorescent phalloidin and imaged. Representative examples of zoospores (DIC: grey) and phalloidin stained actin structures (inverted, black) with an overlay of the two (actin, green) after treatment with each drug. Examples of pseudopods and spikes are highlighted, note pseudopods in SMIFH2 treated cells (highlighted by an arrowhead labeled P) are rounder and less protrusive. Pseudopods (P), actin spikes (S), cortical actin (Co), actin patches (Pa). Scale bar, 10 μm.

**Figure S7.**
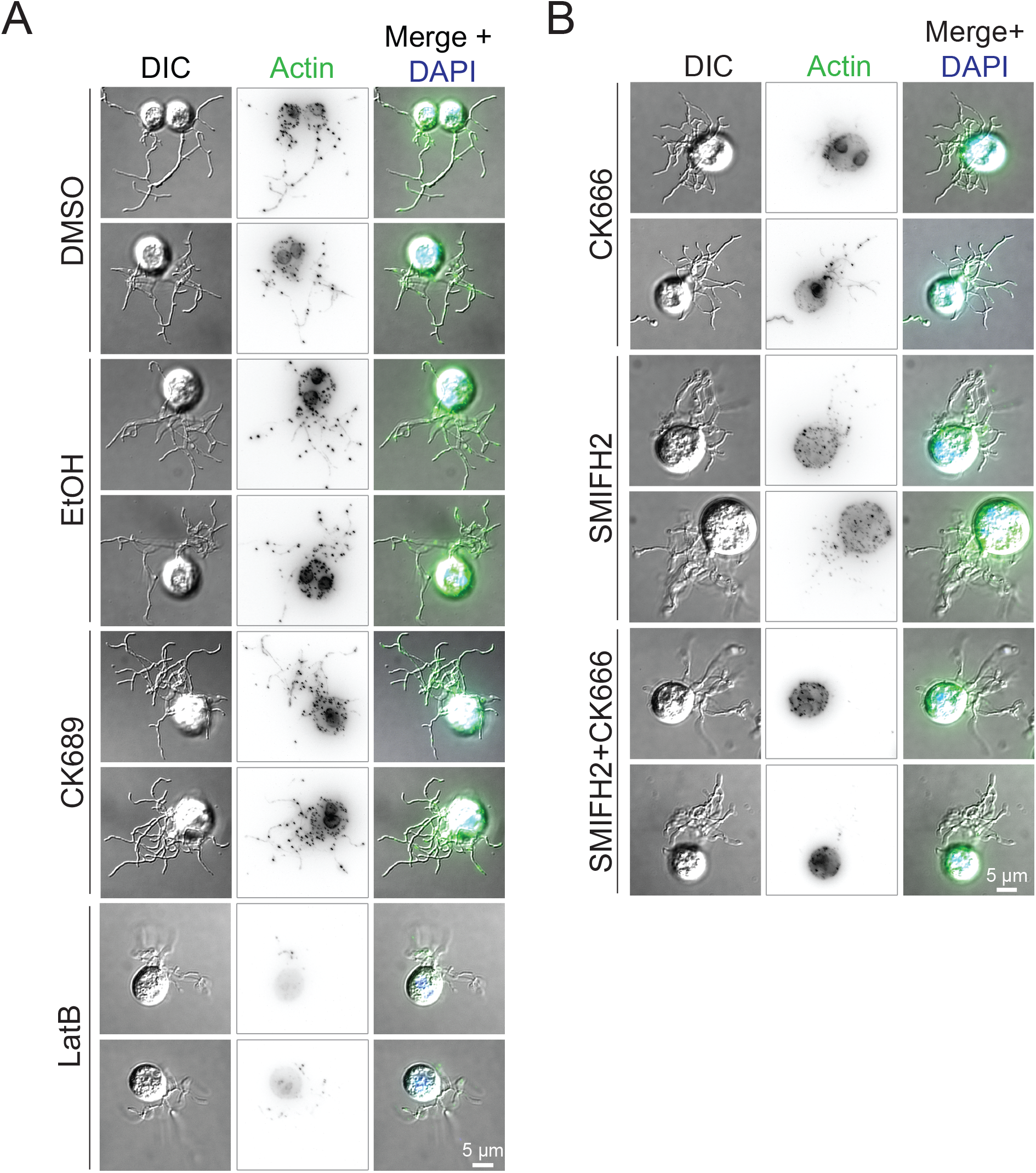
Additional examples of actin structures in Bd sporangia. Populations of *Bd* sporangia seeded 1 day prior were treated with drugs using the same concentrations as in **Figure 6**, then fixed and stained for polymerized actin with phalloidin and for DNA with DAPI. (A-B) Examples of sporangia (DIC: grey) and phalloidin stained actin patches (alone inverted, black; overlay, green), with an overlay including the nucleus (blue) after treatment with each drug. Scale bar, 5 μm.

***Table S1. Accession numbers for chytrid, budding and fission yeast, human, Dictyostelium, and Arabidopsis actin and actin regulatory proteins***. The genbank/NCBI accession numbers for the homologs of actin and actin regulatory proteins in the given chytrid species. Chytrid homologs were identified using BLAST with queries from budding and fission yeast, humans, *Dictyostelium discoideum*, and *Arabidopsis thaliana*; uniprot accession numbers given.

***Table S2. Accession numbers for chytrid, budding and fission yeast, human, Dictyostelium, and Arabidopsis myosin motor proteins***. The genbank/NCBI and Uniprot accession numbers for the homologs of myosin motor proteins in the given chytrid species and budding and fission yeast, humans, *Dictyostelium discoideum*, and *Arabidopsis thaliana*.

## SUPPLEMENTAL DATA FILES

***Supplemental Data File 1. Full alignment of actin and actin related proteins from chytrids, yeast, animals, and other organisms.*** Chytrid sequences sharing ≥50% sequence identity with actin from rabbit, budding yeast, fission yeast, or *Dictyostelium* were compiled and aligned with these actin sequences as well as actin sequences from *Giardia intestinalis (GIAIN)* and *Chlamydomonas reinhardtii* (*CHLRE)*; Arp1 from human, budding and fission yeast, and *Dictyostelium*; and Arp2 and Arp3 from human, budding and fission yeast, *Dictyostelium* and the five chytrids. TCoffee was used to align the sequences (default parameters). The alignment was not edited, and was used for Maximum Likelihood phylogeny estimation (see **Fig. S3**). The beginnings of NCBI/Genbank accession numbers for chytrid sequences are as follows: XP_0066, *Batrachochytrium dendrobatidis (Bd)* JAM81; OAJ, *Bd* JEL423; OON, *Batrachochytrium salamandrivorans*; XP_0166, *Spizellomyces punctatus*; ORY, *Rhizoclosmatium globosum*; KNE, *Allomyces macrogynus*. Non-chytrid sequences are labeled with the Uniprot accession number (if applicable) and the protein and organism name except for Arp2 in *Saccharomyces cerevisiae* (“ARP2”) and *Schizosaccharomyces pombe* (“SPAC11H11.06”). For chytrid actin accession numbers see **Figure S3** and **Supplemental Table 1**.

***Supplementary Data File 2. Alignments of the C-termini of chytrid formins***. The C-terminus of each chytrid formin was hand clipped directly after the end of the FH2 domain, and aligned using TCoffee (default parameters). Sequences with obvious insertions in the diaphanous autoregulatory domain (DAD) region (for reference: starting at XP_016609770.1 M58 (M1240 in the full protein), or position 350 in the alignment) were removed and remaining sequences were realigned as before and evaluated for the presence of the consensus sequences for a DAD (MDXLLXXL; (Alberts 2001; Higgs 2005).

***Supplementary Data File 3. Alignment of formin FH2 domains from 28 species across eukaryotic taxa***. FH2-domain-containing sequences were obtained from the species distribution sunburst of the FH2 domain page (PF02181) on the Pfam website (v32; pfam.xfam.org; (El-Gebali et al. 2019). Sequences which were ≥98% identical were removed to reduce redundancy and the remaining sequences were aligned to the full HMM for the FH2 domain from Pfam. The FH2 domains were isolated by trimming the alignment according to the FH2 domain of the *Saccharomyces cerevisiae* Bni1p sequence (starting PHKKLKQ; ending ADFINEY), which had been previously hand clipped and provided an accurate judgement for the placement of other FH2 domains. Columns which had <80% occupancy were removed from the alignment. This file was used to create the main formin phylogeny in this paper (see **Fig. 4** and **Supplementary Data File 4**).

***Supplementary Data File 4. Full formin phylogeny tree file***. The raw newick file for the formin phylogeny in **Figure 4**. Built by maximum likelihood method through IQtree webserver (default parameters; (Trifinopoulos et al. 2016)) using the alignment in **Supplementary Data File 3**.

***Supplementary Data File 5. Key to the identification system for proteins used in the formin alignment and phylogeny***. Protein identifiers given by Pfam upon sequence download were renamed for functionality before continuing analysis. This file contains the information used for protein identifiers in **Figure 4**, **Figure 5**, **Supplementary Data File 3 and Supplementary Data File 4**, as well as some additional information. The “PFAM_name” column indicates the original identifier of the sequence when downloaded. The “UNIPROT_ACC” column indicates the uniprot accession number for the sequence. The “organism_name” column gives the name of the species the sequence is from, “Orgnaism_code” gives the species’ Uniprot 5-letter species code, “Organism_strain” indicates the specific strain of the organism the sequence is from. The “Gene_name” column indicates the name of the protein used in the phylogeny. The “ORF_range” indicates the amino acid position of the FH2 domain in the full protein sequence.

***Supplementary Data File 6. Raw quantification of actin phenotypes in Bd zoospores and analyzed data***. The raw data from scoring actin phenotypes in *Batrachochytrium dendrobatidis* zoospores. Pa, patches; S, spike; P, pseudopod; C, cortex. NP, nondescript protrusion. Note: NP was used in our preliminary study design using lower resolution imaging methods. The final data sets were higher resolution and allowed for the distinction between pseudopods and spikes in all cases, resulting in zero “NP” cells. “Uninterpretable” cells are cells for which we were not able to accurately assess phenotypes because they were either clumped in groups, partially out of the frame, or out of focus; these cells were excluded from the analysis. “Haze” indicates cells that had no distinct actin structures, but were positive for phalloidin staining.

