## Supplemental Text 1 for "The actin networks of chytrid fungi reveal evolutionary loss of cytoskeletal complexity in the fungal kingdom"

### *Chytrids have an actin regulatory network resembling that of both animals and fungi*

Because the actin structures in *Bd* resemble actin networks found in both animal cells and in Dikarya, we sought to identify putative actin network components to explore how these structures may be built and regulated. We performed BLAST searches for actin, 43 actin regulatory proteins, and myosin motors across five chytrid species: *Bd*, *Batrachochytrium salamandrivorans* (*Bsal*), *Spizellomyces punctatus* (*Sp*), *Rhizoclosmatium globosum* (*Rg*), and *Allomyces macrogynus* (*Am*) (**Fig. 2, Fig. S3 Fig. S4, Fig. S5, Table S1**), and compared these to homologs from humans, *Arabidopsis thaliana* (*At*), *Dictyostelium discoideum* (*Dd*), *Schizosaccharomyces pombe* (*Spom*), and *Saccharomyces cerevisiae* (*Sc*). These analyses revealed several classes of actin cytoskeletal elements conserved across chytrid species:

<u>*Profilin-actin*</u>: All chytrids encode one canonical actin that shares at least ~87% identity with actin from *Dictyostelium*, and budding and fission yeast, as well as rabbit muscle actin (**Fig. S3a**). *Rg* has two additional actin sequences that share ~83% and ~70% identity with known actins (**Fig. S3a**). All canonical chytrid actin sequences, including the three in *Rg*, form a single clade with 74% bootstrap support with known actin sequences (**Fig. S3b, Data S1**). Chytrid protein sequences sharing ~50-70% similarity to known actin sequences form a separate, highly supported clade (99% bootstrap value) with centractin/ARP1, and likely are not divergent actin sequences (**Fig. S3b, Data S1**).

Most eukaryotes tightly control actin polymerization through binding of actin monomers to profilin. This allows profilin to sequester actin monomers and prevent spontaneous nucleation (Carlsson et al. 1977). Binding to profilin also increases the nucleotide exchange rate of ADP-actin to ATP-actin allowing for interaction of ATP-profilin-actin with barbed ends of filaments (Goldschmidt-Clermont et al. 1992). The genomes of all five chytrid species encode two copies of profilin, each between ~24-54% identical to budding yeast PFY1 (**Fig. 2**). This number is intermediate to the number of profilin genes in Dikarya (1) and other eukaryotes like animals, plants, and amoebae (3-5).

<u>*Actin nucleators*</u>: Another level of actin regulation involves controlling actin nucleation. Spontaneous nucleation of actin into filaments is not thermodynamically favored, and cells use nucleators to induce actin polymerization. Three classes of nucleators are known: the Arp2/3 complex, formins, and tandem actin monomer-binding proteins of nucleation. The Arp2/3 complex is mainly responsible for nucleating branched actin filaments off the side of existing filaments. Chytrids encode a single copy of each member of the Arp2/3 complex, with one exception: *Am* encodes two copies of all Arp2/3 complex members except for ARPC4/Arc19 (**Fig. S4**), similar to larger numbers of other genes (**Fig. 2**) and consistent with a whole genome duplication event in this lineage. The Arp2/3 complex is tightly controlled by many proteins, ensuring it is localized properly and activated appropriately. Several of these Arp2/3 complex activators are conserved in humans, chytrids, and Dikarya, such as WISH/Sip1/SPIN90 (with the exception of *Bsal*), and WASP (**Fig. S4**). Only one Arp2/3 complex activator, the SCAR/WAVE complex, is present in humans and most chytrids, but not other fungi. The only chytrid to not have homologs for the SCAR/WAVE complex is *Rg* (**Fig. S4**). A few Arp2/3 complex activators have been lost in all fungi including chytrids: WASH complex and Cortactin (**Fig. S4**).

The second main class of actin nucleators are the formin proteins that nucleate unbranched actin filaments *de novo* and can elongate filaments through processive addition of actin monomers to the growing end (Pruyne et al. 2002; Sagot et al. 2002; Kovar et al. 2003; Zigmond et al. 2003; Moseley et al. 2004; Romero et al. 2004; Kovar et al. 2006). Formins are highly diverse in terms of domain organization, cellular functions, and number of formin genes per species. The conserved element between all formins is the formin homology 2 (FH2) domain that is both necessary and sufficient for actin nucleation (Pruyne et al. 2002; Sagot et al. 2002; Kovar et al. 2003; Li and Higgs 2003). Using a combination of PSI-BLAST and pfam, we found that chytrids have between 4 and 9 formin proteins: *Bd* JAM81 and *Sp* each have 4 FH2-domain-containing sequences; Bd JEL423, *Rg,* and *Bs* each have 5; and *Am* has 9 (**Fig. 2, Table S1**). Some full formin sequences spanned two or more gene annotations (**Table S1**). Chytrid formins have a diversity of domain organizations outside of the FH2 domain, which we discuss below.

The final class of actin nucleators is the tandem actin monomer-binding proteins of nucleation that recruit multiple actin monomers using Wiskott-Aldrich homology 2 (WH2) domains to form an actin nucleus (Quinlan et al. 2005; Ahuja et al. 2007; Rebowski et al. 2008). This class includes SPIRE, which works synergistically with formins in establishing polarity in oocytes and has yet to be found outside of the metazoans (Quinlan et al. 2005; Bradley et al. 2020). Unsurprisingly, a SPIRE homolog was not found in chytrids (**Fig. 2**).

*Capping proteins and actin severing proteins*: Actin is also subject to negative regulation, particularly by capping proteins that prevent further filament elongation and severing proteins that cut existing filaments. Many of these proteins are conserved in humans, chytrids, and Dikarya, such as primary capping proteins CapZ and AIP1, primary severing proteins Colfilin and Twinfilin, as well as the severing catalyst SRV2 (**Fig. 2**). The only protein/protein family to be conserved in animals and chytrids, but not in other fungi, is the Gelsolin/villin family of proteins (**Fig. 2**). Chytrids encode several gelsolin-domain-containing proteins, but relationships to individual members of the Gelsolin/villin family remains to be investigated (**Fig. 2**). It is important to note that plants have 5 villin-like proteins and a gelsolin-domain containing protein that are phylogenetically distinct from other eukaryotic members of this family (Ghoshdastider et al. 2013). The capping proteins tropomodulin and EPS8 are lost in all fungi (**Fig. 2**).

*Actin binding proteins and other actin regulators:* Finally, the remaining actin regulatory proteins we searched for are the actin binding proteins, including proteins involved in endo-/exocytosis, as well as a few other actin regulators which did not neatly fit into another category. Many proteins in this group have been conserved in humans, chytrids, and Dikarya, including: Verprolin/WIP (with the exception of *Rg* and *Am*); Tropomyosin; Fimbrin/plastin; α-actinin (with the exception of *Rg* and *Sc*); endo-/exocytosis proteins EPS15/Ede1, HIP1R/Sla2, and drebrin-like/ABP1 (with the exception of *Sp* for drebrin-like/ABP1), and the Arp2/3 recruiter Coronin (**Fig. 2**). The only protein conserved in humans and chytrids, but not Dikarya is talin (with the exception of *Rg*), which creates the link between adhesion receptors and the actin cytoskeleton (**Fig. 2**). The actin-binding proteins lost in fungi include: Thymosin β4, ENA/VASP family proteins, Missing-in-metastasis (MIM), EPLIN/LIMA1, Espin, Fascin, filamin, and the capping protein regulator CARMIL (**Fig. 2**).

*Myosin motor proteins:* The myosin superfamily of actin-based motors plays diverse roles in cells, including providing contractile forces during cell migration and cytokinesis, powering organelle transport, driving endocytosis and building or maintaining actin-based structures such as filopodia. Myosins all share a motor domain that contains ATP and actin binding sites followed by a lever arm that amplifies the powerstroke and is the site of light chain binding and then a tail region. A small number of myosin family have N terminal extensions that either encompass protein binding modules (e.g. Myo15, Myo18) or an enzyme (e.g. Myo3). The kinetics of a given myosin motor are tuned to its cellular function (processive transporter, ensemble force generation, anchor) and the tail region has a major role in targeting a given myosin to its cellular location (Bloemink and Geeves 2011; Masters et al. 2017). A comprehensive cataloging of myosins shows that chytrids have the same core group of myosins seen in the majority of fungi - Myo1, Myo2, Myo5 and Myo17 (**Fig. S5**) with some species also having Myo22 (Odronitz and Kollmar 2007; Kollmar and Mühlhausen 2017).

Myo1s are ancient, widely expressed myosins that link membranes to the actin cytoskeleton (McIntosh and Ostap 2016; Kollmar and Mühlhausen 2017). These myosins are low duty ratio motors (they are bound to actin for only a short time during their ATPase cycle) meaning that a team of these motors collaborates to execute their cellular function. The Myo1s have conserved roles in endocytosis in fungi, amoebae and mammalian cells where they are critical for efficient internalization and fission of endocytic vesicles. They are also implicated in psuedopod formation and adhesion in migratory cells (McIntosh and Ostap 2016). All fungi, including the chytrids, only express the 'amoeboid' or 'long-tailed' subclass of Myo1s with tails with a membrane binding region that targets these myosins to acidic phospholipids, a domain rich in Gly Pro and Ala that binds actin and an SH3 domain. *Bd*, *Bsal* and *Rg* have a single Myo1, while *Sp* and *Am* have two Myo1s that likely have overlapping functions. Yeast and amoebae Myo1s are regulated by phosphorylation of a conserved Ser/Thr in the motor domain by a PAK kinase, the so-called TEDS site, and the chytrid Myo1s all have this conserved site suggesting that their activities are also regulated by phosphorylation. Unlike the Myo1s from budding and fission yeast, the chytrid Myo1 tails lack a C-terminal acidic region. The budding and fission yeast Myo1s are localized to actin-rich cortical patches that are sites of endocytosis. There they recruit activators of the Arp2/3 complex that interact with their SH3 domain, contributing to the targeted growth of an actin network and focussing actin polymerization forces against the membrane vesicle as it matures, driving internalization (Giblin et al. 2011). The chytrid Myo1s are likely to be localized to the cortical actin patches observed in *Bd* and *Sp* zoospores and sporangia (**Fig 1**; (Medina et al. 2020)).

The Myo2s are found in Amoebozoans and Opisthokonts where they are an essential component of the cytokinetic contractile ring, generating forces necessary for the scission of two daughter cells during the final steps of mitosis (West-Foyle and Robinson 2012). Myo2s are also found in Heteroloboseans, but their function in these species remains unexplored (Fritz-Laylin et al. 2010). Myo2s also play roles in adhesion, morphogenesis, they drive the retrograde flow of the actin network and generate cell polarity in migrating cells by contracting the actin network at the cell rear (Aguilar-Cuenca et al. 2014). Myo2s are low duty ratio motors and work as an ensemble, assembling into bipolar thick filaments via association of their long a-helical coiled tail regions. Unlike muscle myosins, these filaments are dynamic and are only assembled when and where needed (i.e. during cell division). The chytrid fungi have a single Myo2 (**Fig. S5**), as is typical for most fungi, although some species such as *S. pombe* have two. In addition to a role in cytokinesis, budding yeast Myo2 has been implicated in maintaining cell wall integrity by an as yet unknown mechanism (Rodriguez and Paterson 1990).

Myo5s are present in Amoebozoa, Apusozoa and Opisthokonts where they have roles as acitn-based transporters and also act to localize their cargo to the actin-rich network underlying that plasma membrane (Titus 2018). Myo5s have a long lever arm and a correspondingly large powerstroke (i.e they take big steps along actin, 36 nm on average) with a tail that consists of a region of coiled coil that promotes dimerization and a globular C-terminus that binds cargo. Myo5s can be dimeric high duty ratio motors (i.e. they are bound to actin for the majority of their ATPase cycle) or monomeric low duty ratio motors that are organized into ensembles by binding partners, enabling them to function as transporters (Hammer and Sellers 2011). Budding and fission yeast lack the interphase microtubule network typical of other eukaryotic cells (amoebae, mammalian cells) and instead use actin cables for Myo5-dependent distribution of organelles and RNPs. Hyphal fungi use microtubules for long-distance transport and in these species Myo5s work in collaboration with kinesins to transport cargo and also to maintain localization at the cortex (e.g the growing hyphal tip). Many fungi have two Myo5s that perform distinctive functions in cells. For example, *Sc* Myo5 is required for organelle inheritance, polarized budding and mitotic spindle orientation while *Sc* Myo4 is critical for polarized localization of cell fate determinants (Matsui 2003). *Bd*, *Bsal* and *Sp* have a single Myo5 while two are present in *Am* and *Sp* and they likely to play critical roles in intracellular transport, aiding in organelle segregation during division or targeting vesicles to sites of polarized growth.

The Myo17s are unusual chimeric fungal myosins with a core motor domain that lacks a lever arm fused to a cytochrome b5 heme/steroid binding domain then a chitin synthase 2 (Ch2) domain (Kollmar and Mühlhausen 2017). Transmembrane domains in their C-termini orient the motor domain into the cytoplasm and synthase domain inside vesicles that contain this myosin, meaning that upon exocytosis the chitin synthase enzyme is positioned in the outer cell wall. Interestingly, a highly similar myosin is also found in molluscs (Weiss et al. 2006). Myo17s are essential for cell wall integrity and virulence in several pathogenic fungi (Takeshita et al. 2006; Gandía et al. 2014). The *Ustilago maydis* Myo17, Msc1, is found in vesicles that are transported along both microtubules and actin cables by the coordinated action of a kinesin and a Myo5 to the growing hyphal tip (Schuster et al. 2012). The isolated motor domain of Myo17 exhibits ATP-sensitive binding to actin, a hallmark of myosins, and although motor activity is required for Msc1 localization and function, it does not move actin filaments in vitro (Takeshita et al. 2005; Treitschke et al. 2010; Schuster et al. 2012). This suggests that the myosin domain is used to tether the Msc1 to the actin cortex once at the hyphal tip. Chytrid fungi have multiple, highly similar Myo17 family members, as is seen for other fungi - *Am* has 11 Myo17s, *Bd* and *Bs* have 5, *Sp* has 4 and *Rg* has 3. The abundance of these myosins in chytrid fungi is consistent with a likely need for targeted synthesis of their cell wall during the different stages of growth.

The Myo22s are a member of a widely expressed group of myosins with one or two MyTH-FERM domains in the C-terminal tail region (Kollmar and Mühlhausen 2017). These myosins are largely associated with the formation and function of cellular protrusions composed of parallel bundles of actin, such as filopodia and they have also been implicated in cellular adhesion (Tuxworth et al. 2001; Petersen et al. 2016; Weck et al. 2017). Surprisingly, a small group of chytrid fungi including *Rg* and *Sp* have a single Myo22. Neither *Am*, *Bd* nor *Bsal* have a Myo22 and it is not found in any other fungal species outside of the chytrids, revealing that this myosin was lost early in the evolution of fungi. The potential function of Myo22 in *Rg* and *Sp* is unclear at present but it may contribute to the formation of a distinctive actin-based structure in those species that is not present in *Am*, *Bd* or *Bs*.

Taken together, we find chytrid genomes encode a network of actin cytoskeletal elements that is intermediate to the networks of animals and fungi, including SCAR/WAVE complex, gelsolin/villin family proteins, and talin, which are all found in animals and chytrids but not Dikarya.

**Chytrid formins have additional domain architectures beyond diaphanous-like and PTEN-like**

*Non-typical diaphanous-like formins:* In addition to the typical diaphanous-related formins, *Am* has two proteins that are likely diaphanous-related, but each uniquely differs from the archetypal structure: one is missing a recognizable diaphanous autoregulatory domain (Genbank: KNE63115.1; Uniprot: A0A0L0SL59); and the other has no predicted FH3 domain and has a potentially functional diaphanous autoregulatory domain (Genbank: KNE56028.1; Uniprot: A0A0L0S0G4) (**Fig. 3; Data S2**). *Bd, Bs,* and *Rg* each have a diaphanous-like formin that lacks a GTPase-binding domain (**Fig. 3**). In *Rg* this formin (Genbank: ORY46026.1; Uniprot: A0A1Y2CGH6) also lacks an FH1-domain and has a truncated C-terminus (**Fig 3**). There are potentially functional diaphanous autoregulatory domains present in the *Bd* (NCBI Reference sequence: XP_006675776.1; Uniprot F4NSE8) and *Bs* (Genbank: OON09123.1; Uniprot: A0A1S8W533) sequences (**Fig. 3, Data S2**).

A unique diaphanous-like formin domain architecture has been identified in *Rg* (ORY50340.1), which contains an ApaG domain (PF04379), a GTPase-binding domain, an FH3 and a very short FH1 domain, all N-terminal to the FH2 domain (**Fig. 3**). This domain is not known to be in formin proteins and although the function of the ApaG domain is currently unknown, proteins with ApaG domains include the ApaG proteins in bacteria and some F-box proteins in mammals (Gibson et al. 1991; Ilyin et al. 2000).

*Other formin architectures:* *Bs* and *Am* each have two formins containing an N-terminal FH2 domain, typical of the inverted formins mammalian subfamily (**Fig. 3**). However, unlike other inverted formins (Hegsted et al. 2017), these formins have no obvious FH1 domain, although one of these formins in *Bs* (Genbank: OON02955.1; Uniprot: A0A1S8VKL4) has a potentially functional diaphanous autoregulatory domain (**Fig. 3, Data S2**).
